# Activated Ras/JNK driven Dilp8 in imaginal discs adversely affects organismal homeostasis during early pupal stage in *Drosophila*, a new checkpoint for development

**DOI:** 10.1101/049882

**Authors:** Mukulika Ray, Subhash C. Lakhotia

**Affiliations:** Cytogenetics Laboratory, Department of Zoology, Banaras Hindu University, India 221005

**Author notes:** Author for correspondence, Email: S. C. Lakhotia, Mukulika Ray, Tele.

**Keywords:** Halloween genes, Ptth, Ecdysone, *hsrω* lncRNA, inter-organ signaling

## Abstract

**Background:** Dilp8-mediated inhibition of ecdysone synthesis and pupation in holometabolous insects maintains developmental homeostasis through stringent control of timing and strength of moulting signals. We examined reasons for normal pupation but early pupal death observed in certain cases.

**Results:** Over-expression of activated Ras in developing eye/wing discs inhibited Ptth expression in brain via up-regulated JNK signaling mediated Dilp8 secretion from imaginal discs, which inhibited ecdysone synthesis in prothoracic gland *after* pupariation, leading to death of ~25-30 Hr old pupae. Inhibition of elevated Ras signaling completely rescued early pupal death while post-pupation administration of ecdysone to organisms with elevated Ras signaling in eye discs partially rescued their early pupal death. Unlike the earlier known Dilp8 action in delaying pupation, hyperactivated Ras mediated elevation of pJNK signaling in imaginal discs caused Dilp8 secretion *after* pupariation. Ectopic expression of certain other transgene causing pupal lethality similarly enhanced pJNK and early pupal Dilp8. Sub-optimal ecdysone levels after 8 Hr of pupation prevented the early pupal metamorphic changes and caused organismal death.

**Conclusions:** Our results reveal early pupal stage as a novel Dilp8 mediated post-pupariation checkpoint and provide further evidence for inter-organ signaling during development, wherein a peripheral tissue influences the CNS driven endocrine function.

## Introduction

Regulation of signaling pathways during development of an organism is essential for normal development. In order to maintain the overall homeostasis for proper running of organism level biological processes, different signaling pathways are modulated in specific cell types of a given tissue at specific time and to a regulated extent. Any change in fine tuning of the spatial and temporal, qualitative as well as quantitative, activation of signaling molecule/s hampers proper biological functioning through cascading affects. Pervasiveness of this effect has become more evident with increasing understanding of inter-organ signaling (Rajan and Perrimon, 2011; Droujinine and Perrimon, 2013; Droujinine and Perrimon, 2016; Sinclair et al., 2017).

During our earlier studies on interaction between Ras signaling and *hsrω* lncRNAs, we found that the ectopically induced high levels of Ras signaling cascade in eye discs of *Drosophila* third instar larvae culminated in pupal death (Ray et al., 2019b). This study showed that while the *sev-GAL4* driven ectopic expression of activated Ras in developing eye discs led to substantial late pupal death as pharates, still higher levels of activated Ras in eye discs, achieved by co-expression of *sev-GAL4* driven activated Ras and *hsrω-RNAi* caused early pupal death (~25-30 Hr after pupation). Pupal death following ectopic expression of certain transgenes driven by predominantly eye-specific *sev-GAL4* or *GMR-GAL4* drivers has been reported earlier as well (Morris et al., 2006; Mallik and Lakhotia, 2009b). Since eyes are not essential for survival of flies, the observed pupal death in all these instances has been rather intriguing. We have examined possible reasons for the early pupal death following induction of very high levels of activated Ras in larval imaginal discs.

Recent studies have shown that disruption in optimal larval development affects ecdysone synthesis and consequently delays pupation via signaling through Dilp8, an insulin like peptide (Colombani et al., 2012; Garelli et al., 2012; Colombani et al., 2015; Garelli et al., 2015; Okamoto and Yamanaka, 2015). Unlike these earlier results on role of Dilp8 in delaying pupation, we found that very high Ras/JNK activity, which disrupts normal development of imaginal discs, enhanced secretion of Dilp8 from eye discs at early pupal stage. Ectopic expression of activated Ras or certain other transgenes like *UAS-rpr* or *UAS-tak^1^* in wing discs also elevated Dilp8 levels in early pupae and resulted in their death. High Dilp8 reduced prothoracicotrophic hormone (Ptth) production in brain, leading to reduced activity of genes involved in ecdysone biosynthesis in prothoracic gland, the seat of ecdysone hormone production (Rewitz et al., 2009; Niwa and Niwa, 2016). Insufficient systemic ecdysone hormone during the critical early pupal stage hampered normal development, causing breakdown of organismal homeostasis and consequent death. These results thus unravel the early pupal ecdysone surge (Handler, 1982) as a new checkpoint for Dilp8 action and explain the intriguing results noted in several other studies (Morris et al., 2006; Mallik and Lakhotia, 2009b) that expression of certain transgenes by predominantly eye-disc specific GAL4 drivers is associated with pupal lethality. An early version of this study was published as a pre-print (Ray and Lakhotia, 2016).

This study shows that elevated Ras signaling can also contribute to Dilp8 expression and how activated Ras signaling in a particular tissue has far reaching consequences on distant organs. It also highlights susceptibility of certain stages of development to specific changes in signaling events, particularly that of Dilp8 on pupal developmental processes and its importance as messenger between epithelia and brain. Since activated Ras is also implicated in various cancers (Karim and Rubin, 1998; Prober and Edgar, 2000; Fernández-Medarde and Santos, 2011; Pylayeva-Gupta et al., 2011), the present finding of the activated Ras dependent long-range inter-organ signaling has implications for disease conditions as well.

## Materials and Methods

### Fly stocks

All fly stocks and crosses were maintained on standard agar cornmeal medium at 24±1^0^C. The following stocks were obtained from the Bloomington Stock Centre (USA): *w^1118^; sev-GAL4*; + (no. 5793), *w^1118^; UAS-GFP* (no. 1521), *w^1118^; UAS-rpr* (no. 5824), *w^1118^; UAS-Tak1* (no. 58810), *ecd^1^* (no. 218) and *w^1118^; UAS-Ras1^V12^* (no. 4847). The other stocks, viz., *w^1118^; GMR-GAL4* (Freeman, 1996), *w^1118^; UAS-hsrω-RNAi^3^* (Mallik and Lakhotia, 2009b), *w^1118^; GMR-GAL4; UAS-hsrω-RNAi^3^, w^1118^; Sp/CyO; dco^2^ e/TM6B* and *w^1118^; UAS-127Q* (Kazemi-Esfarjani and Benzer, 2000), were available in the laboratory. The *UAS-hsrω-RNAi^3^* is a transgenic line used for down regulating the *hsrω*-nuclear transcripts under a GAL4 driver (Mallik and Lakhotia, 2009b). The *UAS-RafRBDFLAG* stock (Freeman et al., 2010) was provided by Dr. S Sanyal (Emory University, USA). Using these stocks, appropriate crosses were made to generate following stocks:

a. w^1118^; sev-GAL4 UAS-GFP; dco^2^ e/TM6B
b. w^1118^; sev-GAL4 UAS-GFP; UAS-hsrω-RNAi^3^
c. w^1118^; UAS-GFP; UAS-Ras1^V12^
d. w^1118^; sev-GAL4 UAS-GFP; ecd^1^
e. w^1118^; UAS-RafRBDFLAG; UAS-Ras1^V12^/TM6B

Some of these were used directly or were further crossed to obtain progenies of following genotypes as required:

a. w^1118^; sev-GAL4 UAS-GFP/UAS-GFP; dco^2^ e/+
b. w^1118^; sev-GAL4 UAS-GFP/UAS-GFP; dco^2^ e/UAS-Ras1^V12^
c. w^1118^; sev-GAL4 UAS-GFP/UAS-GFP; UAS^-^hsrω-RNA^i3/^UAS-Ras1^V12^
d. w^1118^; sev-GAL4 UAS-GFP/UAS-RafRBDFLAG; dco^2^ e/+
e. w^1118^; sev-GAL4 UAS-GFP/UAS-RafRBDFLAG; dco^2^ e/UAS-Ras1^V12^
f. w^1118^; sev-GAL4 UAS-GFP/UAS-RafRBDFLAG; UAS-hsrω-RNAi^3^/UAS-Ras1^V12^
g. w^1118^; GMR-GAL4/UAS-GFP
h. w^1118^; GMR-GAL4/UAS-GFP; +/ UAS-Ras1^V12^
i. w^1118^; GMR-GAL4/UAS-GFP; UAS-hsrω-RNAi^3^/ UAS-Ras1^V12^
j. w^1118^; GMR-GAL4/UAS-rpr
k. w^1118^; GMR-GAL4/UAS-Tak1
l. w^1118^; GMR-GAL4/UAS-127Q.
m. w^1118^ MS1096-GAL4; GMR-GAL4/+
n. w^1118^MS1096-GAL4; GMR-GAL4/UAS-GFP; +/ UAS-Ras1^V12^
o. w^1118^ MS1096-GAL4; +/UAS-rpr
p. w^1118^ MS1096-GAL4; +/UAS-Tak1
q. w^1118^ MS1096-GAL4; +/UAS-127Q

The *w^1118^*, *dco^2^ e* and *UAS-GFP* markers are not mentioned while writing genotypes in Results while the *UAS-hsrω-RNAi^3^* transgene (Mallik and Lakhotia, 2009b) is referred to as *hsrω-RNAi*.

For conditional inhibition of ecdysone synthesis using the temperature-sensitive *ecd^1^* mutant allele (Henrich et al., 1987), the freshly formed *sev-GAL4 UAS-GFP; ecd^1^* pupae, reared from egg-laying till pupation at 24±1^0^C, were transferred to 30±1^0^C for further development.

Flies of desired genotypes were crossed and their progeny eggs were collected at hourly intervals. Larvae that hatched during a period of 1 Hr were separated to obtain synchronously growing population. Likewise, larvae that began pupation during an interval of 1 Hr were separated to obtain pupae of defined age (expressed as Hr after pupa formation or Hr APF).

### GFP imaging

Actively moving late third instar larvae and pupae of desired ages were dissected in Poels’ salt solution (PSS) (Tapadia and Lakhotia, 1997) and tissues fixed in freshly prepared 4% paraformaldehyde (PFA) in phosphate-buffered saline (PBS, 130 mm NaCl, 7 mm Na_2_HPO_4_, 3mm KH_2_PO_4_,pH 7.2) for 20 min. After three 10 min washes in 0.1% PBST (PBS + 0.1% Triton-X-100), the tissues were counterstained with DAPI (4’, 6-diamidino-2-phenylindole dihydrochloride, 1μg/ml) and mounted in 1,4-Diazabicyclo [2.2.2] octane (DABCO) antifade mountant for imaging the *sev-GAL4* or *GMR-GAL4* driven *UAS-GFP* expression.

### Whole organ immunostaining

Brain and eye discs from *sev-GAL4 UAS-GFP/UAS-GFP; dco^2^e*/+, *sev-GAL4 UAS-GFP/UAS-GFP; dco^2^ e/UAS-Ras1^V12^*, *sev-GAL4 UAS-GFP/UAS-GFP* and *sev-GAL4 UAS-GFP/UAS-GFP;UAS-hsrω-RNAi/UAS-Ras1^V12^* actively migrating late third instar larvae and pupae of desired age were dissected in PSS and immediately fixed in freshly prepared 4% paraformaldehyde in PBS for 20 min and processed for immunostaining as described earlier (Prasanth et al., 2000). Rabbit monoclonal anti-pJNK (1:100; Promega) or rat anti-Dilp8 (1:50; gifted by Dr. P. Léopold, France) (Colombani et al., 2012) were used as primary antibodies. Appropriate secondary antibodies conjugated either with Cy3 (1:200, Sigma-Aldrich, India) or Alexa Fluor 633 (1:200; Molecular Probes, USA) or Alexa Fluor 546 (1:200; Molecular Probes, USA) were used to detect the given primary antibody. Chromatin was counterstained with DAPI (1μg/ml). Tissues were mounted in DABCO for confocal microscopy with Zeiss LSM Meta 510 using Plan-Apo 40X (1.3-NA) or 63X (1.4-NA) oil immersion objectives. Quantitative estimates of the proteins in different regions of eye discs were obtained with the help of Histo option of the Zeiss LSM Meta 510 software as described earlier (Ray et al., 2019b). All images were assembled using the Adobe Photoshop CS3 software.

### Measurement of ecdysone levels

Fifteen pupae of the desired stages and genotypes were collected in 1.5 ml tubes and stored at −70°C in methanol till further processing. They were homogenized in methanol and centrifuged at 13000 rpm following which the pellets were re-extracted in ethanol and air dried (Ou et al., 2011). The dried extracts were dissolved in EIA (**E**nzyme **I**mmuno**a**ssay) buffer at 4°C overnight prior to the enzyme immunoassay. The 20E-EIA antiserum (#482202), 20E AChE tracer (#482200), precoated (Mouse Anti-Rabbit IgG) EIA 96-Well Plates (#400007), and Ellman’s Reagent (#400050) were obtained from Cayman Chemical (USA), and assays were performed according to the manufacturer’s instructions.

### Exposure of pupae to external ecdysone

Fresh 0-1 Hr old *ecd^1^* white prepupae, grown through the larval period at 24°C, were transferred to 30°C to inhibit further ecdysone synthesis. After 8 Hr at 30°C, they were transferred to ecdysone solution (1μg/ml water) for 12 Hr at 30°C following which they were removed from ecdysone solution and allowed to develop at 30°C. Similarly, 8-9 Hr old *sev-GAL4>Ras1^V12^hsrω*-RNAi pupae were incubated in ecdysone solution (1μg/ml) for 12 Hr and allowed to develop further at 24°C. For each genotype, parallel controls were maintained without the external ecdysone treatment. Numbers of pupae dying at early (25 - 30 Hr APF) or late stages was scored in each case.

### Microarray Analysis

RNA was isolated from 16-17 Hr old *sev-GAL4>UAS-GFP, sev-GAL4>Ras1^V12^* and *sev-GAL4>Ras1^V12^ hsrω-RNAi* pupae using TriReagent (Sigma-Aldrich) as per manufacturer’s instructions. Microarray analysis of these RNA samples was performed on Affymetrix Drosophila Genome 2.0 microarray chips for 3’ IVT array following the Affymetrix GeneChip Expression Analysis Technical manual using the GeneChip 3’ IVT Plus Reagent Kit, Affymetrix GeneChip^®^ Fluidics station 450, GeneChip^®^ Hybridization oven 645 and GeneChip^®^Scanner 3000. Summary of the expression levels for each gene in the four genotypes was obtained from the Affymetrix Transcription analysis console and was subjected to Gene ontology search using David Bioinformatics software (https://david.ncifcrf.gov). The microarray data have been deposited at GEO (http://www.ncbi.nlm.nih.gov/geo/query/acc.cgi?acc=GSE80703).

### Real Time Quantitative Reverse transcription–PCR (RT-qPCR)

Total RNAs were isolated from eye discs and brain ganglia of late third instar larvae and early pupal stages of the desired genotypes using TriReagent as per the manufacturer’s (Sigma-Aldrich) instructions. First-strand cDNAs were synthesized as described earlier (Mallik and Lakhotia, 2009b). The prepared cDNAs were subjected to real time PCR using forward and reverse primer pairs listed in Table 1. Real time qPCR was performed using 5μl qPCR Master Mix (Syber Green, Thermo Scientific), 2 picomol/μl of each primer per reaction in 10 μl of final volume in ABI 7500 Real time PCR machine.

**Table 1.**
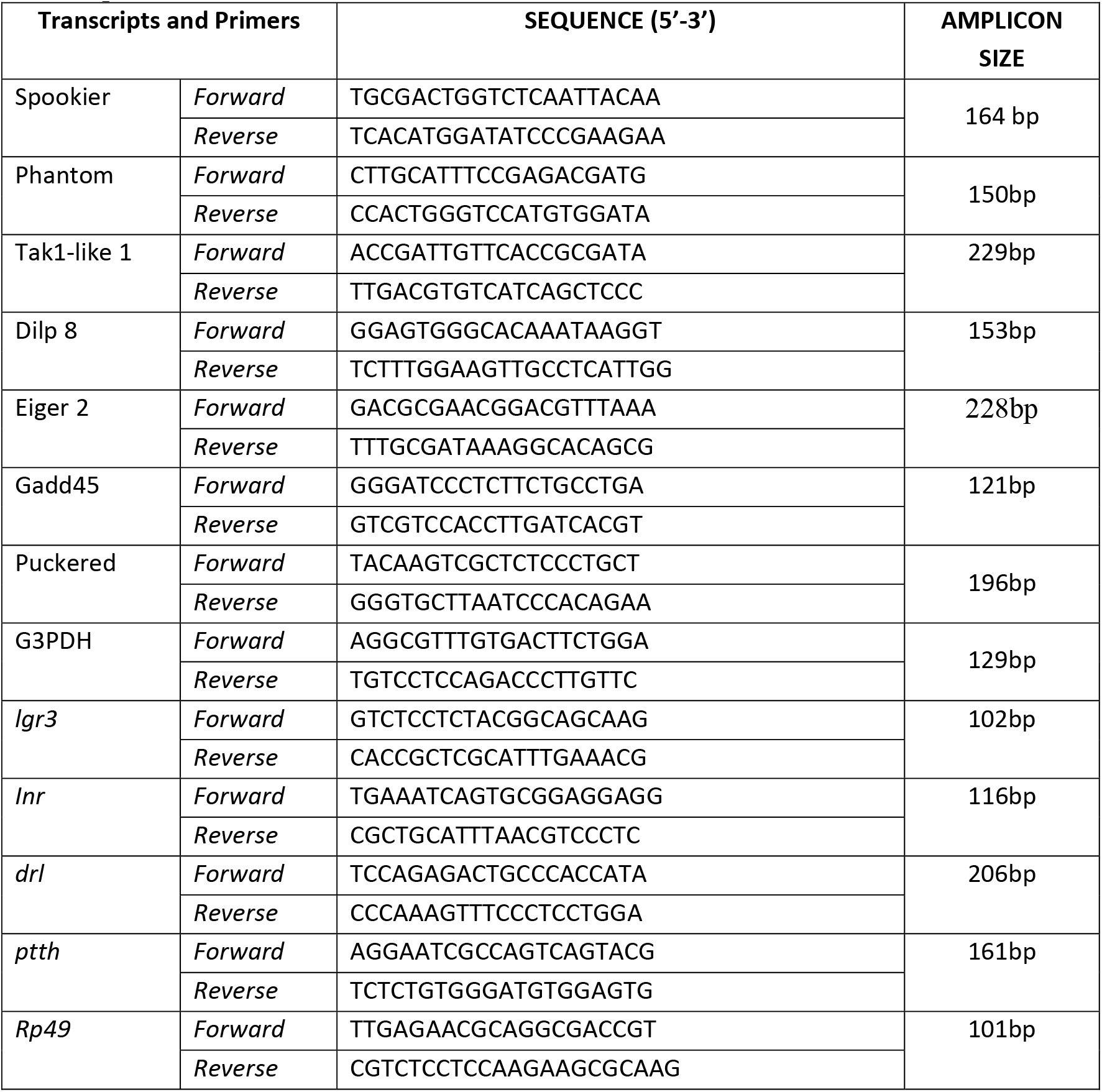
Forward and reverse primers used for qRT-PCR for quantification of different transcripts

## Results

### Enhanced Ras activity in eye discs results in defects in pupal development and early to late pupal lethality depending upon degree of Ras activity

*UAS-Ras1^V12^* transgene expression has been widely used for sustained Ras signaling since the Ras1^V12^ protein does not require upstream events for activation (Karim et al., 1996). In the present study we expressed the activated Ras1^V12^ protein in eye disc cells using *GMR-GAL4* or *sev-GAL4* drivers, the former driving expression in a much greater number of cells in eye discs than the later (Ray and Lakhotia, 2015), so that the Ras signaling is much more widespread in eye discs expressing the *GMR-GAL4* driver. All the *GMR-GAL4>UAS-Ras1^V12^* individuals died at early pupal stage (between 25-30 Hr APF), whereas the *sev-GAL4>UAS-Ras1^V12^* pupae developed normally till late pupal stages but a majority (~88%, N=1058) died as pharates while the rest eclosed with rough eyes due to extra R7 photoreceptors (Ray et al., 2019b). Up-regulation of Ras activity further in *sev-GAL4* expressing eye discs (Ray et al., 2019b) by co-expressing *hsrω-RNAi* transgene (Mallik and Lakhotia, 2009b) that down regulates the *hsrω* lncRNAs, resulted in early pupal death of all *sev-GAL4>UAS-Ras1^V12^ hsrω-RNAi* like that in the *GMR-GAL4>UAS-Ras^V12^*. Normally, the pupae undergo head eversion after 12 Hr of pupation so that the 12-14 Hr APF stage wild type pupae show a proper demarcation of head, thorax and abdomen (Fig 1A). All *sev-GAL4>UAS-Ras1^V12^* pupae also showed similar demarcation of body divisions (Fig 1B). However, in the case of *GMR-GAL4>UAS-Ras1^V12^* and *sev-GAL4>UAS-Ras1^V12^ hsrω-RNAi* pupae such demarcation was completely absent (Fig 1C, D). Further, the Malpighian tubules (MT), which become abdominal by 12-14 Hr APF in wild type as well as *sev-GAL4>UAS-Ras1^V12^* pupae (arrows in Fig. 1A, B), did not retract but persisted in anterior part in *GMR-GAL4>UAS-Ras1^V12^* and *sev-GAL4>UAS-Ras1^V12^ hsrω-RNAi* pupae (arrows in Fig 1C, D). The characteristic metamorphic changes causing compaction of cells in MT (Fig. 1E), histolysis of SG (Fig. 1G) and disappearance of the posterior 6 pairs of segmental dorso-median *sev-GAL4>GFP* expressing neurons (Ray and Lakhotia, 2015) (Fig. 1I, K) seen in wild type and *sev-GAL4>UAS-Ras1^V12^* 24-25 Hr old pupae, did not occur in the early dying 24-25 Hr old *sev-GAL4>UAS-Ras1^V12^ hsrω-RNAi* pupae. They continued to show these structures as in 8-9 Hr old pupae (Fig. 1F, H, J, L). Likewise, the *GMR-GAL4>UAS-Ras1^V12^* at 24-25 Hr pupae carried intact SG while the MT did not acquire the characteristic compact cell arrangement (Fig. 1M, N). The *GMR-GAL4>GFP* expressing dorso-median paired neurons in ventral ganglia also persisted in 24-25 Hr old *GMR-GAL4>UAS-Ras1^V12^* pupae (not shown). It is significant that absence of metamorphic changes during development of pupa and the early pupal death (~25-30 Hr APF) in *GMR-GAL4>UAS-Ras1^V12^* and *sev-GAL4>UAS-Ras1^V12^ hsrω-RNAi* genotypes correlates with the earlier reported (Ray et al., 2019b) wider and stronger Ras signaling in their eye discs than in the *sev-GAL4>UAS-Ras1^V12^* eye discs (Fig. 1O).

**Fig 1.**
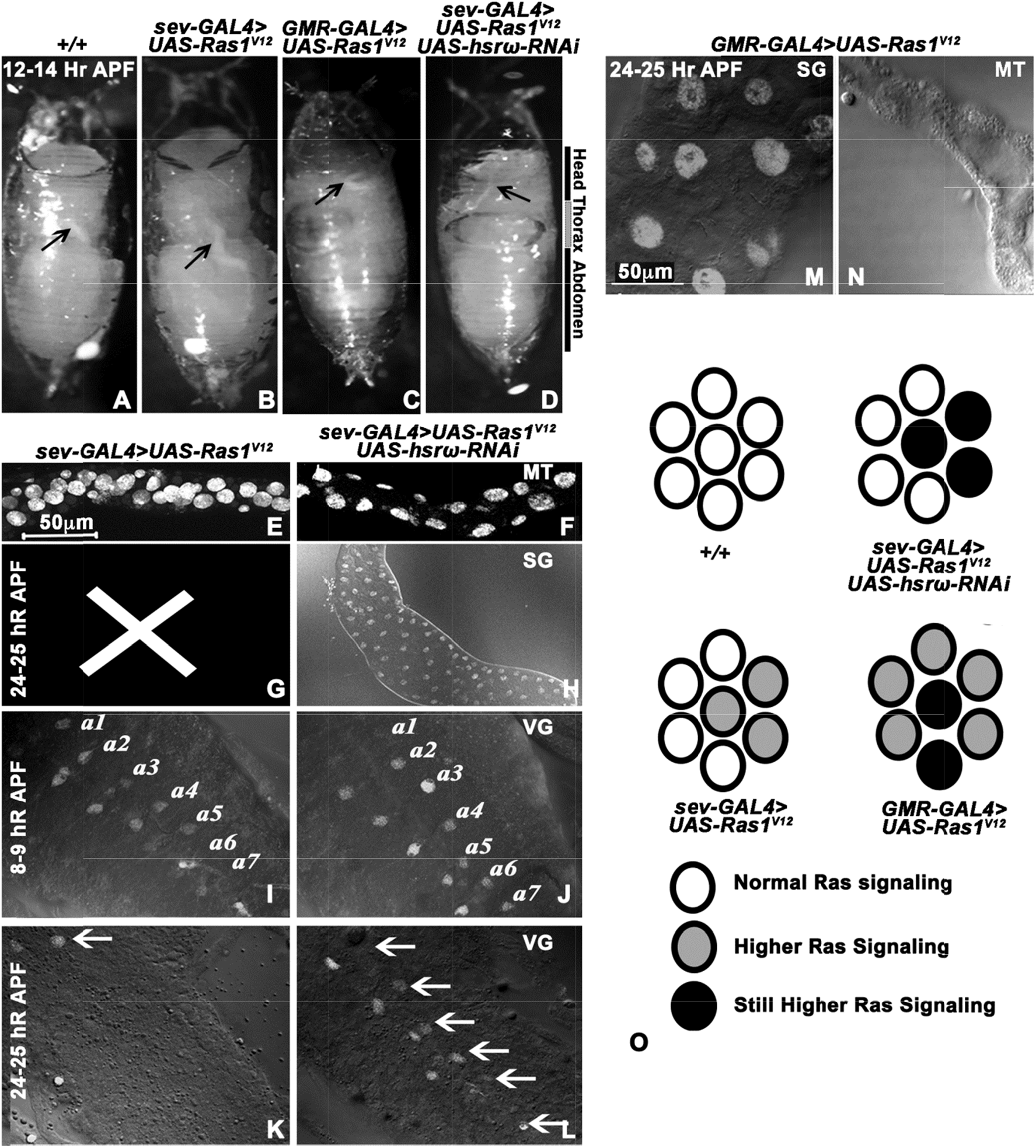
The prepupal to pupal metamorphic changes are affected in pupae that die early following highly enhanced Ras signaling in eye discs. **A**-**D** Photomicrographs of 12-14 Hr old pupae of genotypes noted on top of each; relative positions of head, thorax and abdomen are marked on the right while arrows on the images point to the distal white segment of the anterior Malpighian tubules. **E**-**H** Confocal and DIC images of parts of DAPI stained (white) Malpighian tubules (MT, **E, F**) and salivary glands (SG, **G, H**) in 24-25 Hr old *sev-GAL4>UAS-Ras1^V12^* (**E, G**) and *sev-GAL4>UAS-Ras1^V12^ hsrω-RNAi* (**F, H**) pupae; a cross is marked in **G** since the SG is already histolysed and thus not visible in *sev-GAL4>UAS-Ras^V12^* pupae at this stage. **I**-**L** Confocal and DIC images of parts of ventral ganglia showing *sev-GAL4>GFP* (white) expression in a1-a7 segmentally arranged dorso-median pairs of neurons at 8-9 Hr (**I, J**) and 24-25 Hr (**K, L**) APF stage in genotypes noted above the columns; arrows in **K and L** indicate the persisting *sev-GAL4>GFP* expressing dorso-median paired neurons. Scale bars in **E** and **G** denote 50μm and apply to **E-F** and **G-J**, respectively. **M**-**N** Confocal and DIC images of parts of DAPI stained (white) salivary glands (SG, **M**) and Malpighian tubules (MT, **N**) in 24-25 Hr old *GMR-GAL4>UAS-Ras1^V12^* pupae. **O** Schematics of cells with varying levels of activated Ras in ommatidial units in eye discs of the indicated genotypes (based on (Ray et al., 2019b)).

### Transcriptome analysis of early dying pupae showed down regulation of ecdysone synthesis pathway genes and up regulation of JNK pathway genes

Since eyes are dispensable for survival of *Drosophila* (Sang and Jackson, 2005), the pupal death following over-expression of activated Ras under *sev-GAL4* or *GMR-GAL4* drivers that pre-dominantly express in eye discs (Ray and Lakhotia, 2015) was unexpected. In order to understand the organism level changes that may cause the early pupal death, we carried out microarray based whole transcriptome analysis of 16-17 Hr old *sev-GAL4> UAS-GFP, sev-GAL4>UAS-Ras1^V12^* and *sev-GAL4>UAS-Ras1^V12^ hsrω-RNAi* pupae. As reported earlier (Ray et al., 2019b), these three genotypes have normal, high and very high Ras signaling, respectively (Fig. 1O). A pair wise comparison of expression levels between different genotypes revealed significant up- or down-regulation of many genes, belonging to diverse GO categories, in *sev-GAL4>UAS-Ras1^V12^hsr ω-RNAi* sample (Table 2).

**Table 2.** Major pathways affected in individuals co-expressing *sev-GAL4* driven activated Ras (*Ras1^V12^*) and *hsrω-RNAi* when compared with those expressing *sev-GAL4* driven activated Ras only.

| Pathways down-regulated in *sev-GAL4>UAS-Ras1<sup>V12</sup> hsrω-RNAi* | Pathways up-regulated in *sev-GAL4>UAS-Ras1<sup>V12</sup> hsrω-RNAi* |  |  |
| --- | --- | --- | --- |
| GO:0008610: lipid biosynthetic process<br>GO:0005976: polysaccharide metabolic process<br>GO:0002376: immune system process<br>GO:0045087: innate immune response<br>GO:0006694: steroid biosynthetic process | GO:0006030: chitin metabolic process<br>GO:0006022: aminoglycan metabolic process<br>GO:0016052: carbohydrate catabolic process<br>GO:0005976: polysaccharide metabolic process<br>GO:0006950: response to stress<br>GO:0009266: response to temperature stimulus<br>GO:0009308: amine metabolic process<br>GO:0005975: carbohydrate metabolic process<br>GO:0006629: lipid metabolic process<br>GO:0045087: innate immune response<br>GO:0006955: immune response<br>GO:0008063: Toll signaling pathway<br>GO:0008219: cell death<br>GO:0035070: salivary gland histolysis<br>GO:0048102: autophagic cell death<br>GO:0016271: tissue death<br>GO:0007559: histolysis<br>GO:0012501: programmed cell death<br>GO:0006865: amino acid transport<br>GO:0006508: proteolysis | GO:0006582: melanin metabolic process<br>GO:0006570: tyrosine metabolic process<br>GO:0007160: cell-matrix adhesion<br>GO:0031589: cell-substrate adhesion<br>GO:0007155: cell adhesion<br>GO:0010631: epithelial cell migration<br>GO:0031644: regulation of neurological system process<br>GO:0050890: cognition<br>GO:0022008: neurogenesis<br>GO:0042180: cellular ketone metabolic process<br>GO:0006575: cellular amino acid derivative metabolic process<br>GO:0043436: oxoacid metabolic process<br>GO:0006082: organic acid metabolic process<br>GO:0019752: carboxylic acid metabolic process<br>GO:0019471: 4-hydroxyproline metabolic process<br>GO:0031098: stress-activated protein kinase signaling pathway | GO:0007600: sensory perception<br>GO:0006816: calcium ion transport<br>GO:0007568: aging GO:0007165: signal transduction<br>GO:0007188: G-protein signaling, coupled to cAMP nucleotide second messenger<br>GO:0007187: G-protein signaling, coupled to cyclic nucleotide second messenger<br>GO:0007166: cell surface receptor linked signal transduction<br>GO:0035272: exocrine system development<br>GO:0007254: JNK cascade |

**Note:** The pathways that appear to contribute to the early pupal lethality in *sev-GAL4>UAS-Ras1<sup>V12</sup> hsr ω-RNAi* genotype are highlighted in gray.

Several genes involved in lipid and steroid biosynthesis processes were down-regulated in *sev-GAL4>UAS-Ras1^V12^ hsrω-RNAi* pupae that show early pupal death (Table 3). This was significant since the early dying pupae failed to undergo the metamorphic changes that normally follow the ecdysone pulse beginning at 8-10 Hr APF (Handler, 1982). Three *(spookier, phantom* and *shadow)* of the five 20-hydroxy-ecdysone biosynthesis related Halloween genes (Huang et al., 2008) showed down-regulation in *sev-GAL4>UAS-Ras1^V12^ hsrω-RNAi* when compared with *sev-GAL4>UAS-Ras1^V12^* pupae (Table 3), while *shade* did not change and *disembodied* displayed up-regulation. The *spookier* and *phantom* genes act at early steps in ecdysone biosynthesis (Huang et al., 2008) while the *shadow* gene product converts 2-deoxyecdysone into ecdysone (Warren et al., 2002).

**Table 3.**
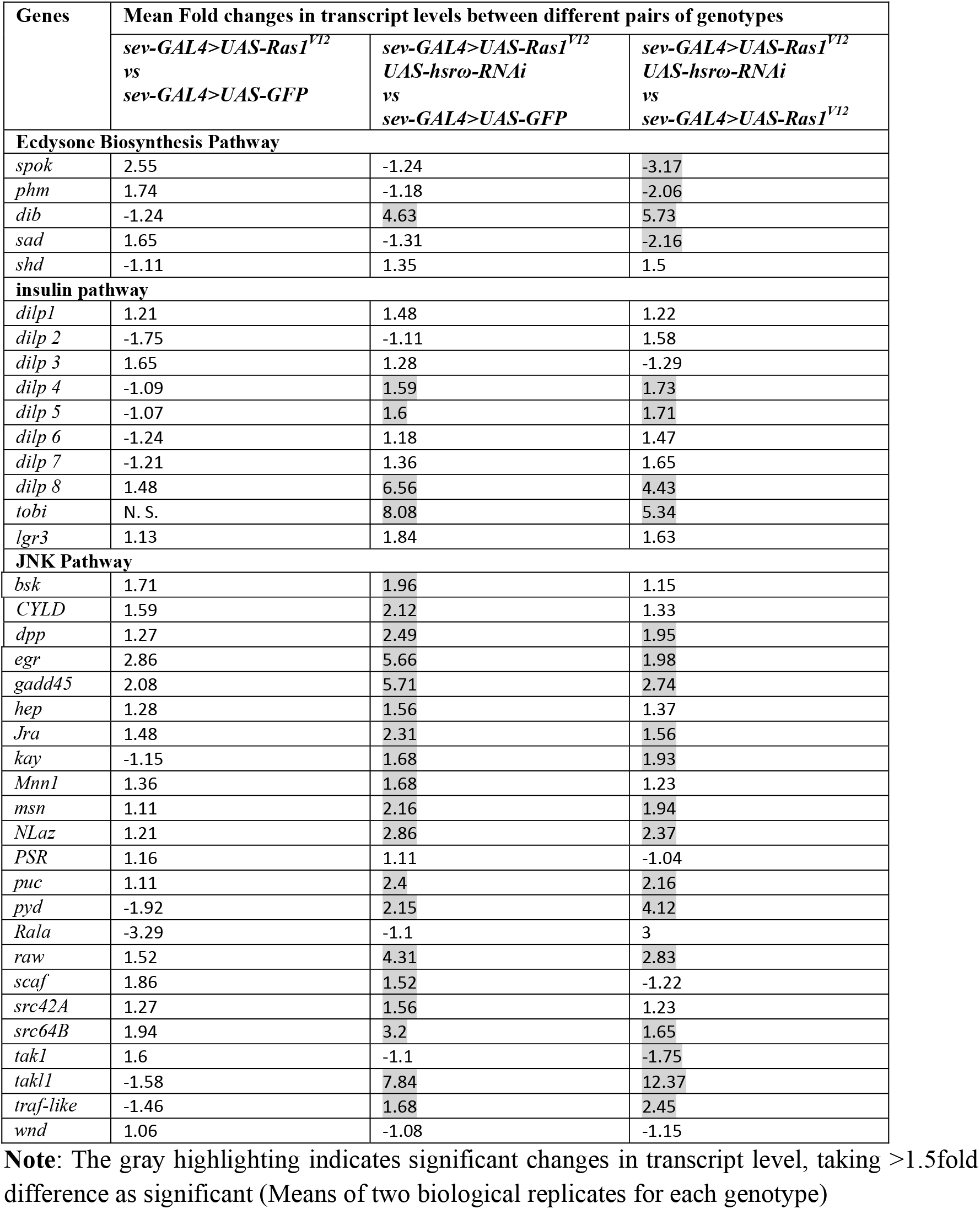
Transcription of several genes in steroid and ecdysone biosynthesis, insulin- and JNK-signaling pathways is affected in early dying pupae with *sev-GAL4* driven enhanced Ras signaling in eye disc cells

The up-regulated groups of genes (Table 2) in *sev-GAL4>UAS-Ras1^V12^ hsrω-RNAi* pupae included, among many others, those associated with stress responses, cell and tissue death pathways, autophagy, toll signaling, innate immune system etc. Some of these seem to be primarily due to activated Ras or altered *hsrω* expression since the Ras/MAPK pathway is known to regulate immune response (Ragab et al., 2011; Hauling et al., 2014) while the *hsrω* transcripts have roles in stress responses, cell death and several other pathways (Mallik and Lakhotia, 2009a; Lakhotia, 2011; Lakhotia, 2016; Ray et al., 2019a; Ray et al., 2019b). However, beside the predicted up-regulation of G-protein coupled signal transduction pathway following ectopic expression of activated Ras, a significant increase in expression of several JNK pathway genes was also noted (Tables 2 and 3).

Damaged or tumorous imaginal discs are reported to affect ecdysone synthesis through up regulation of Dilp8 secretion possibly through the JNK mediated enhanced Dilp8 synthesis (Colombani et al., 2012; Garelli et al., 2012). Of the eight known Dilps (Dilp1-8) in *Drosophila* (Nässel et al., 2013), only *dilp8* showed significant up-regulation in *sev-GAL4>UAS-Ras1^V12^ hsrω-RNAi* when compared with *sev-GAL4>UAS-GFP* or *sev-GAL4>UAS-Ras1^V12^* (Table 3). Like the *dilp8,* transcripts of *tobi (target of brain insulin)* were also significantly up-regulated (Table 3) when *sev-GAL4>UAS-Ras1^V12^* was co-expressed with *hsrω-RNAi*. The Dilp8 action on ecdysone synthesis in prothoracic gland is reported to involve the neuronal relaxin receptor Lgr3 (Colombani et al., 2015; Garelli et al., 2015). Our microarray data for whole body pupal RNA indicated that the *lgr3* transcripts were only marginally elevated in *sev-GAL4>UAS-Ras1^V12^ hsrω-RNAi* expressing early pupae.

These results suggested that the enhanced Ras signaling in *sev-GAL4>UAS-Ras1^V12^ hsrω-RNAi* eye discs may affect ecdysone synthesis in prothoracic gland via up regulation of Dilp8 peptide secretion from the eye discs. Therefore, we checked levels of 20-hydroxy-ecdysone, the functional ecdysone hormone, and validated the results of microarray analysis using qRT-PCR in different larval and pupal tissues (eye disc, prothoracic gland and brain) of *sev-GAL4>UAS-Ras1^V12^* (high Ras activity), *sev-GAL4>UAS-Ras1^V12^ hsrω-RNAi* and *GMR-GAL4>UAS-Ras1^V12^* (very high Ras activity).

### Early dying pupae with enhanced Ras activity in eye discs have reduced levels of post-pupation ecdysone

Enzyme Immunoassay for 20-hydroxy-ecdysone revealed ecdysone levels in 8-9 Hr old pupae expressing *sev-GAL4>UAS-Ras1^V12^* alone or together with *hsrω-RNAi* transgene to be comparable to those in similar age *sev-GAL4>UAS-GFP* pupae (Fig. 2A). However, 24-25 Hr old pupae co-expressing *hsrω-RNAi* with *sev-GAL4>UAS-Ras1^V12^* did not display the characteristic increase in ecdysone levels (Handler, 1982) seen in *sev-GAL4>UAS-GFP* or *sev-GAL4>UAS-Ras1^V12^* (Fig. 2A).

**Fig. 2.**
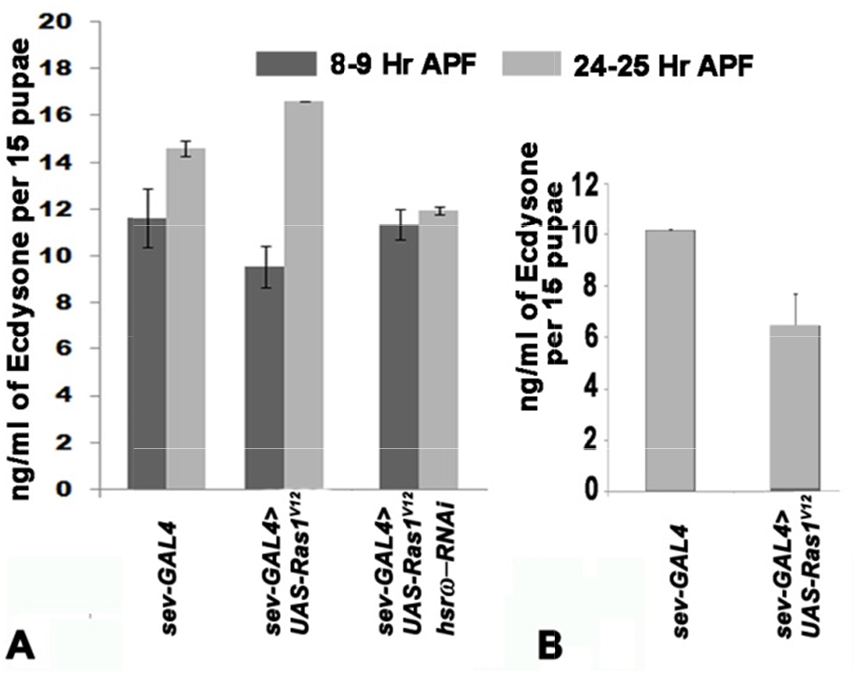
Ectopically induced high Ras signaling in eye discs is associated with reduced levels of systemic 20-hydroxyecdysone. **A-B** Bars showing mean (+S.E. of two independent replicates) levels of 20 hydroxyecdysone (Y axis) in different genotypes (X-axis) at 8-9 Hr APF (dark grey) and at 24-25 Hr APF (light grey). Each genotype also carries *UAS-GFP* transgene.

The ecdysone levels in *GMR-GAL4>UAS-Ras1^V12^* 24-25 Hr old pupae were also significantly lower than in the corresponding age *GMR-GAL4>UAS-GFP* control pupae (Fig 2B).

Further support for reduced levels of ecdysone being responsible for the early pupal death was obtained by examining phenotypes of otherwise wild type early pupae in which the ecdysone levels were conditionally reduced using the temperature sensitive *ecd^1^* allele, which inhibits production of ecdysone when grown at the 29-30°C (Henrich et al., 1987). The *sev-GAL4>UAS-GFP; ecd^1^* larvae were reared since hatching at 24°C and 0-1 Hr old white prepupae were transferred to 30°C to conditionally inhibit further ecdysone synthesis. About 50% of these pupae showed early death by 25-30 Hr APF while the remaining died as late pupae (Table 4). The early dying *sev-GAL4>UAS-GFP; ecd^1^* pupae, reared since beginning of pupation at 30°C, showed persistence of the segmental dorso-median pairs of *sev-GAL4>GFP* expressing neurons (Fig. 3A), less compact arrangement of MT cells (Fig. 3B) and the other developmental defects similar to those displayed by the *sev-GAL4>UAS-Ras1^V12^ hsrω-RNAi* or *GMR-GAL4>UAS-Ras1^V12^* early dying pupae.

**Table 4.**
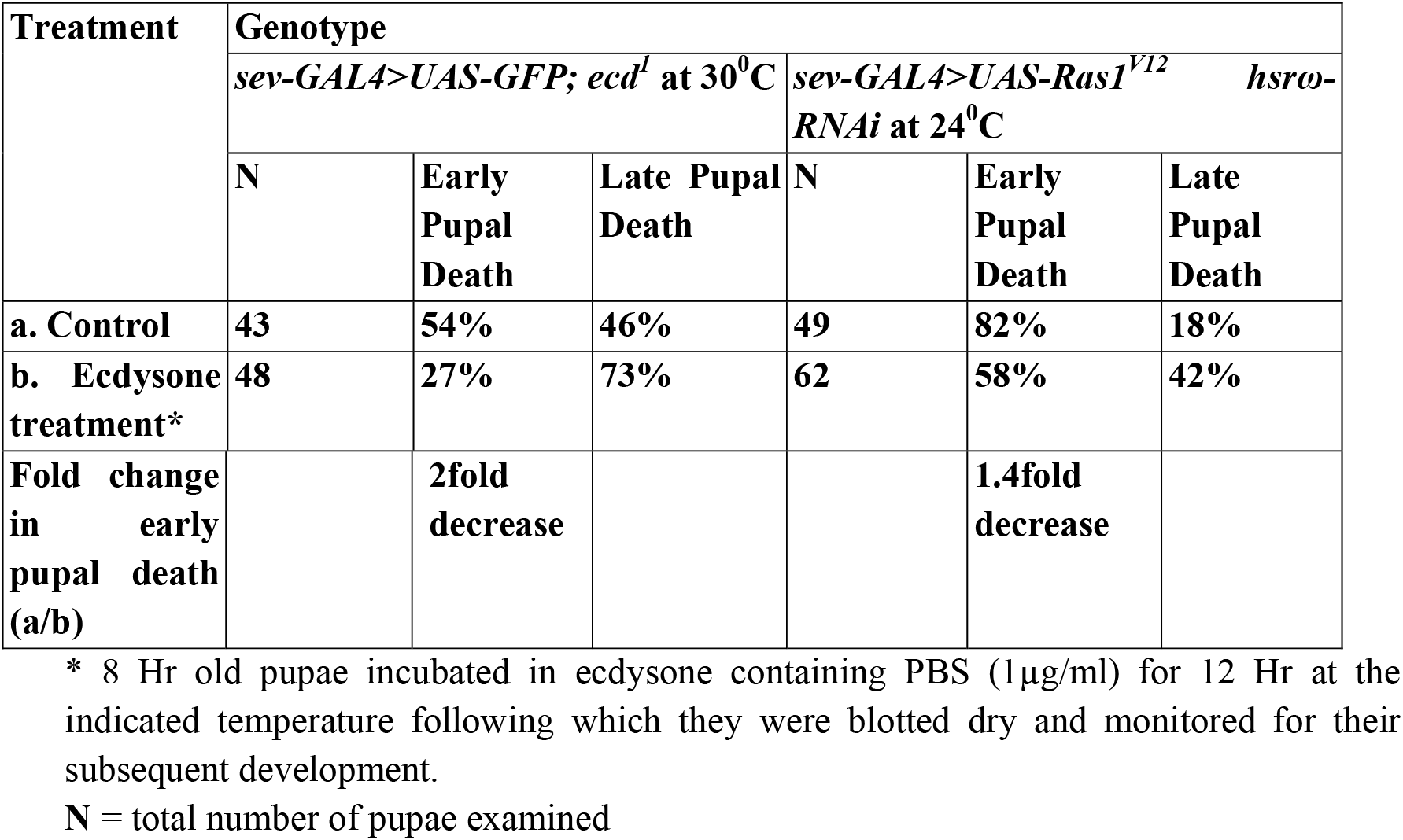
Exogenous ecdysone partially suppressed early pupal death in *sev-GAL4>UAS-GFP; ecd^1^* (maintained at restrictive temperature from 0-1Hr pre-pupa onwards), *sev-GAL4>UAS-Ras1^V12^ hsrω-RNAi* pupae

**Fig. 3.**
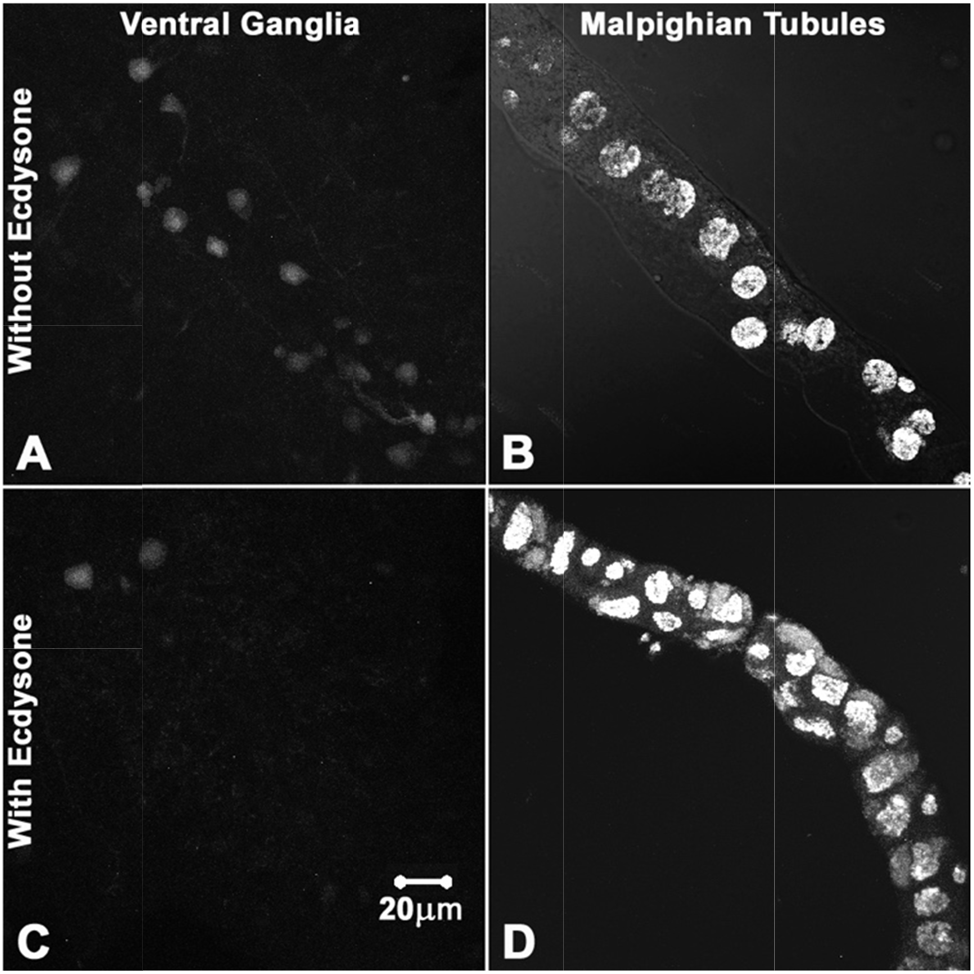
Exogenous ecdysone promotes prepupal to pupal metamorphic changes in *ecd^1^* mutant pupae maintained at the restrictive temperature (30^0^C). **A**, **C** Confocal projections images showing GFP (white) expression in ventral ganglia in 24-25 Hr old *sevGAL4>UAS-GFP; ecd^1^* pupae maintained at 30^0^C since the beginning of pupation without (**A**) or with (**C**) exposure to exogenous ecdysone between 8-20 Hr APF. **B**, **D** Confocal projection images of DAPI-stained (white) Malpighian tubules of 24-25 Hr old *sevGAL4>UAS-GFP; ecd^1^* pupae maintained at 30^0^C since the beginning of pupation without (**B**) or with (**D**) exposure to exogenous ecdysone between 8-20 Hr APF. The scale bar in **C** applies to all the images.

### Externally provided ecdysone partially rescued the early pupal death

We exposed 8-9 Hr old pupae of different genotypes to exogenous ecdysone by incubating them in 1 μg/ml 20-hydroxyecdysone dissolved in PBS for 12 Hr, following which they were taken out, blotted dry and allowed to develop further under normal condition. Incubation in PBS alone did not affect pupal development in any genotype. To confirm effectiveness of such exogenous exposure to ecdysone, we used *ecd^1^* pupae as control. Fresh 0-1 Hr old *sev-GAL4>UAS-GFP; ecd^1^* white prepupae, grown through the larval period at 24°C, were transferred to 30°C to block further ecdysone synthesis. After 8 Hr at 30°C, they were transferred to the 20-hydroxy-ecdysone solution for 12 Hr at 30°C to keep the endogenous ecdysone synthesis inhibited, while it was provided in the incubation medium. After the 12 Hr ecdysone exposure, the pupae were returned to 30°C for further development. As seen from the data in Table 4, about 54% *sev-GAL4>UAS-GFP; ecd^1^* pupae maintained at 30°C from 0-1 Hr pre-pupa stage onwards without any exogenous ecdysone died by 25-30 Hr APF. Significantly, there was a 2-fold reduction in the early dying *ecd^1^* pupae maintained at 30°C but exposed to exogenous ecdysone from 8 to 20 Hr APF (Table 4), although none eclosed. Significantly, the pre-pupal to pupal metamorphic changes in these surviving pupae occurred normally so that the *sev-GAL4* driven GFP expressing 6 pairs of abdominal dorso-median neurons in ventral ganglia disappeared (Fig. 3C), while the MT cells attained the expected compaction in ecdysone treated *ecd^1^* pupae maintained at 30°C (Fig. 3D). Survival of a significant proportion of *ecd^1^* pupae, maintained at 30°C while being exposed to exogenous ecdysone, clearly indicated that the moulting hormone penetrated the early pupal case in at least some of them.

Exposure of 8 Hr old *sev-GAL4>UAS-Ras1^V12^ hsrω-RNAi* pupae to exogenous ecdysone for 12 Hr at 24°C resulted in a 1.4-fold decrease in those dying early, although none of the survivors emerged as flies (Table 4). The somewhat lesser decrease (1.4-fold, Table 4) in death of the ecdysone exposed *sev-GAL4>UAS-Ras1^V12^ hsrω-RNAi* pupae beyond the early stage may be related to the fact that, compared to 30°C reared *ecd^1^* pupae (54%), a larger proportion (82%) of *sev-GAL4>UAS-Ras1^V12^ hsrω-RNAi* pupae died early (Table 4).

It is significant that persistence of the entire set of 7 pairs of these neurons at 24-25 Hr APF stage in *ecd^1^* mutants reared at restrictive temperature since pupation (Fig. 3A) is similar to that in *GMR-GAL4>UAS-Ras1^V12^* and *sev-GAL4>UAS-Ras1^V12^ hsrω-RNAi* genotypes (Fig. 1L). Disappearance of the posterior six pairs of these neurons in the restrictive temperature reared surviving *ecd^1^* and *sev-GAL4>UAS-Ras1^V12^ hsrω-RNAi* pupae following exogenous ecdysone treatment strongly indicates that it is the lack of ecdysone which leads to persistence of GFP expression in these paired neurons till 24-25 Hr APF stage, rather than their being the causal factor for the reduced levels of ecdysone in these genotypes.

Together, these results confirm that *sev-GAL4>UAS-Ras1^V12^ hsrω-RNAi* and *GMR-GAL4>UAS-Ras1^V12^* pupae fail to release optimal levels of ecdysone after the 8-9 Hr pupal stage and therefore, do not undergo the expected metamorphic changes.

### Dilp8 protein levels are elevated in early pupal eye discs in proportion with elevated Ras levels

It is reported that in instances of delayed pupation, ecdysone synthesis is inhibited by Dilp8 signaling (Colombani et al., 2012; Garelli et al., 2012). Therefore, to know if the reduced levels of ecdysone noted above in early dying pupae, we next examined Dilp8 protein levels in eye discs with normal (*sev-GAL4>UAS-GFP*), high (*sev-GAL4>UAS-Ras^V12^*) and very high (*sev-GAL4>UAS-Ras^V12^ hsrω-RNAi*) Ras activity (Ray et al., 2019b). Immunostaining of late third instar larval eye discs with Dilp8 antibody (Colombani et al., 2012) revealed absence of Dilp8 staining in the ommatidial cells (Fig. 4A, B, E, F, I, J) but a faint presence of Dilp8 in cytoplasm of peripodial cells in all the three genotypes (Fig. 4C, D, G, H, K, L). The Dilp8 staining in ommatidial cells of 8-9 Hr pupal eye discs co-expressing *sev-GAL4>UAS-Ras1^V12^* and *hsrω-RNAi* was slightly enhanced (Fig. 4U-V), but their peripodial cells showed much stronger levels (Fig. 3W-X) than their same age counterparts in *sev-GAL4>UAS-GFP* (Fig. 4M-P) or *sev-GAL4>UAS-Ras1^V12^* (Fig. 4Q-T). The stronger Dilp8 immunostaining in *sev-GAL4>UAS-Ras1^V12^* and *hsrω-RNAi* pupal eye discs was confirmed by quantification of Dilp8 fluorescence (Figure 4Y) using the Histo tool of the LSM510 Meta software on projection images of 12 optical sections through the eye discs in each genotype. The increase in Dilp8 levels in early pupal *sev-GAL4>UAS-Ras1^V12^ hsrω-RNAi* eye discs correlates with the much higher levels of activated Ras (Ray et al., 2019b) in these eye discs than in those expressing only *sev-GAL4>UAS-Ras1^V12^* and with their death as early pupae. The non-cell-autonomous increase in Dlip8 protein in peripodial cells appears to be related to accumulation of secreted products in the peripodial cells prior to their release in the hemolymph.

**Fig. 4.**
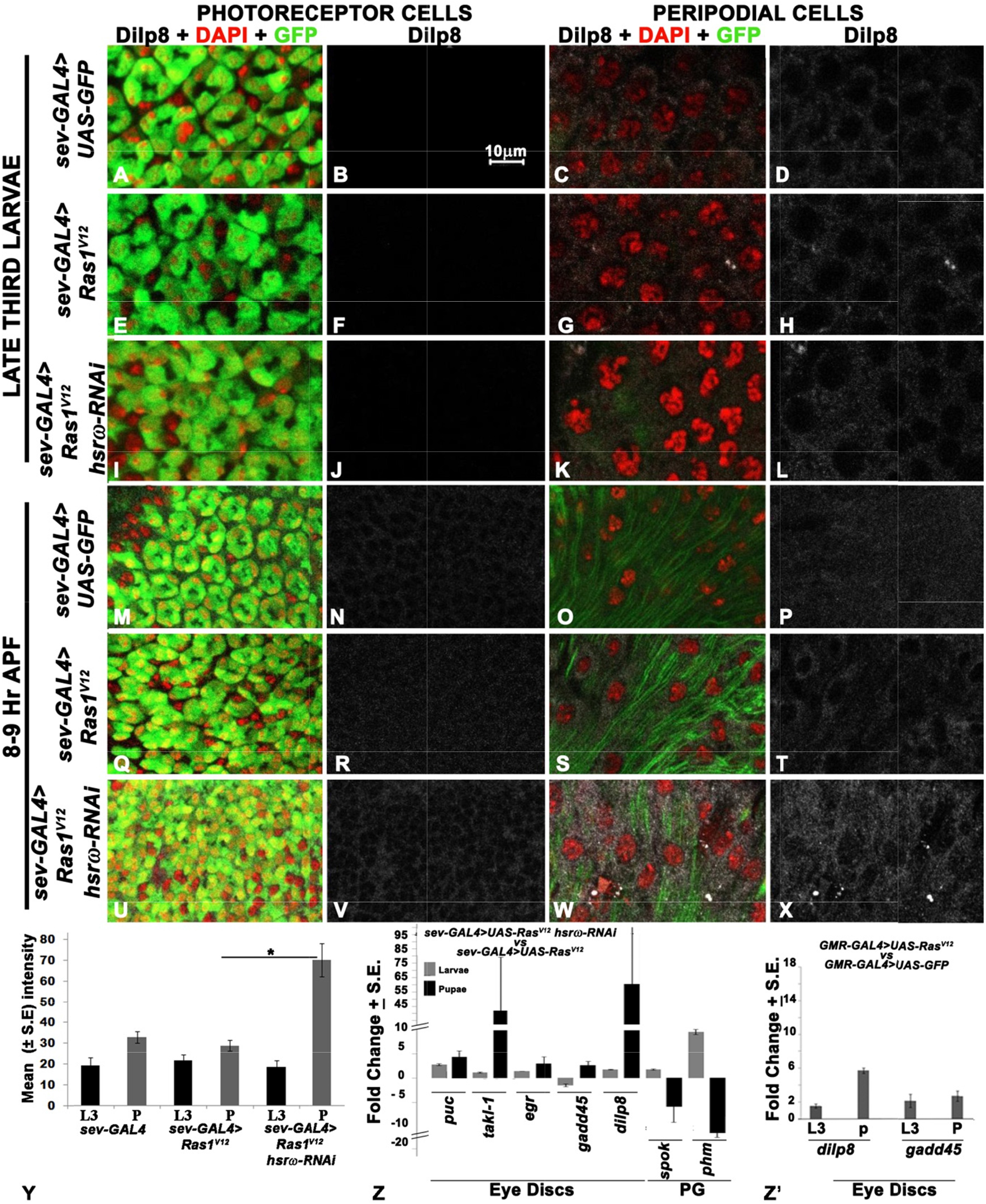
Expression of Dilp8 and several JNK pathway genes remains nearly unchanged in eye discs of third instar larvae but increases in those of 8-9 Hr pupae of genotypes showing early pupal death together with reduction in some of the Haloween gene transcripts in prothoracic gland. **A**-**X** Confocal optical sections of photoreceptor (1st and 2nd columns) and peripodial (3^rd^ and 4^th^ columns) cells in third instar larval (**A**-**L**) and 8-9 Hr pupal eye discs (**M**-**X**), showing Dilp8 (white, 2^nd^ and 4^th^ columns) in different genotypes (noted on left of each row). Combined Dilp8 (white), DAPI (blue) and *GFP* (green) fluorescence confocal images are shown in 1st and 3^rd^ columns. Scale bar in **B** (10μm) applies to all. **Y** Bars showing mean intensity (+S.E., N = 15 eye discs from 3 independent replicates for each genotype) of Dilp8 immunofluorescence (Y axis) in different genotypes (X-axis) in late third instar larval (black, L3) and 8-9 Hr pupal eye discs (grey, P). An asterisk mark above the horizontal line connecting the two compared genotypes indicates significantly (P≤ 0.05) high Dilp8 staning in *sev-GAL4>UAS-Ras1^V12^ UAS-hsrω-RNAi* than in *sev-GAL4>UAS-Ras1^V12^*. **Z** Bars showing qRT-PCR based mean fold changes (+S.E., two biological replicates, Y-axis) in different transcripts in eye discs (ED) or prothoracic glands (PG, X-axis) of *sev-GAL4>UAS-Ras1^V12^ hsrω-RNAi* late 3rd instar larvae (grey bars) or 8-9 Hr old pupae (Black bars) when compared with corresponding *sev-GAL4>UAS-Ras1^V12^* samples. **Z’** Bars showing qRT-PCR based mean fold changes (+S.E., two biological replicates, Y-axis) in levels of *dilp8* and *gadd45* transcripts (X-axis) in eye discs of *GMR-GAL4>UAS-Ras1^V12^* late 3rd instar larvae (L3) or 8-9 Hr old pupae (P) when compared with corresponding *GMR-GAL4>UAS-GFP* samples.

### Increase in Ras signaling in eye discs caused increase in *dilp8* and JNK pathway gene transcripts while Halloween gene activity in prothoracic gland was reduced

We validated some of the microarray data by qRT-PCR using RNA samples from late larval and early pupal (8-9 Hr APF) eye discs and/or prothoracic glands (Fig. 4Z). In agreement with the microarray data for whole pupal RNA, qRT-PCR of 8-9 Hr old pupal eye disc RNA confirmed significant up-regulation of transcripts of the JNK signaling pathway genes like *tak1-like1, eiger, gadd45* and *puckered* in *sev-GAL4>UAS-Ras1^V12^ hsrω-RNAi* when compared with same age *sev-GAL4>UAS-Ras1^V12^* eye discs (Fig. 4Z). Interestingly, late third instar larval eye discs of different genotypes did not show much difference for these transcripts except of the JNK pathway gene *puckered* (Fig. 4Z). However, the increase in *puckered* transcripts in larval eye discs was significantly less than in pupal eye discs.

Quantification of *dilp8* transcripts from late third instar larval and 8-9 Hr pupal eye discs of different genotypes by qRT-PCR revealed, in agreement with microarray data for total pupal RNA, substantial increase (~65 fold) in *dilp8* transcripts in *sev-GAL4>UAS-Ras1^V12^ hsrω-RNAi* pupal eye discs than in those expressing only the activated Ras (Fig. 4Z). However, *dilp8* transcript levels in late third instar larval eye discs showed only a marginal up-regulation in *sev-GAL4>UAS-Ras1^V12^ hsrω-RNAi* eye discs (Fig. 4Z).

Down-regulation of *spookier* and *phantom* transcripts in *sev-GAL4>UAS-Ras1^V12^ hsrω-RNAi* was validated by qRT-PCR using 8-9 Hr old pupal prothoracic gland (PG) RNA. Compared to *sev-GAL4>UAS-Ras1^V12^*, both transcripts showed a high reduction in *sev-GAL4>UAS-Ras1^V12^ hsrω-RNAi* pupal PG (Fig. 4Z). Intriguingly, *sev-GAL4>UAS-Ras1^V12^ hsrω-RNAi* larval prothoracic glands showed up-regulation of *spookier* and *phantom* transcripts (Fig. 4Z).

Levels of *dilp8* and *gadd45* transcripts were elevated in late 3^rd^ instar larval and 8-9 Hr old *GMR-GAL4>UAS-Ras1^V12^* pupal eye discs when compared with corresponding age *GMR-GAL4>UAS-GFP* control eye discs. Interestingly, however, the increase in each case was more pronounced in pupal than in third instar larval eye discs (Fig 4Z’).

### Phosphorylated JNK is elevated in larval and pupal eye discs in early dying genotypes with high Ras signaling in eye discs

Since the tumorous or damaged tissues are known to up-regulate stress responsive JNK pathway (Beira and Paro, 2016) and earlier studies (Colombani et al., 2012; Garelli et al., 2012; Colombani et al., 2015) showed that the JNK pathway plays a key role in Dilp8 synthesis, we checked levels of phosphorylated JNK in third instar larval and 8-9 Hr old pupal eye discs in different genotypes. Immunostaining for phosphorylated JNK (pJNK, Basket) in *sev-GAL4>UAS-GFP* late third instar eye discs revealed low pJNK in photoreceptor cells (Fig. 5A). Photoreceptor cells from *sev-GAL4>UAS-Ras1^V12^* larval discs showed slightly elevated pJNK staining (Fig. 5B). Interestingly, co-expression of *hsrω-RNAi* transgene with *sev-GAL4>UAS-Ras1^V12^* resulted in a much greater increase in pJNK (Fig. 5C) than in *sev-GAL4>UAS-Ras1^V12^* larval eye discs (Fig. 5B). Quantitative data for pJNK immunostaining of larval eye discs, obtained using the Histo option of LSM510 Meta software, presented in Fig. 5D confirm the greatly enhancement in pJNK in *sev-GAL4>UAS-Ras1^V12^ hsrω-RNAi* eye discs (genotype 3 in Fig. 5D). Immunostaining of 8-9 Hr pupal *sev-GAL4>UAS-Ras1^V12^ hsrω-RNAi* eye discs also showed elevated pJNK than in *sev-GAL4> UAS-Ras1^V12^* (Fig. 5E-F).

**Fig. 5.**
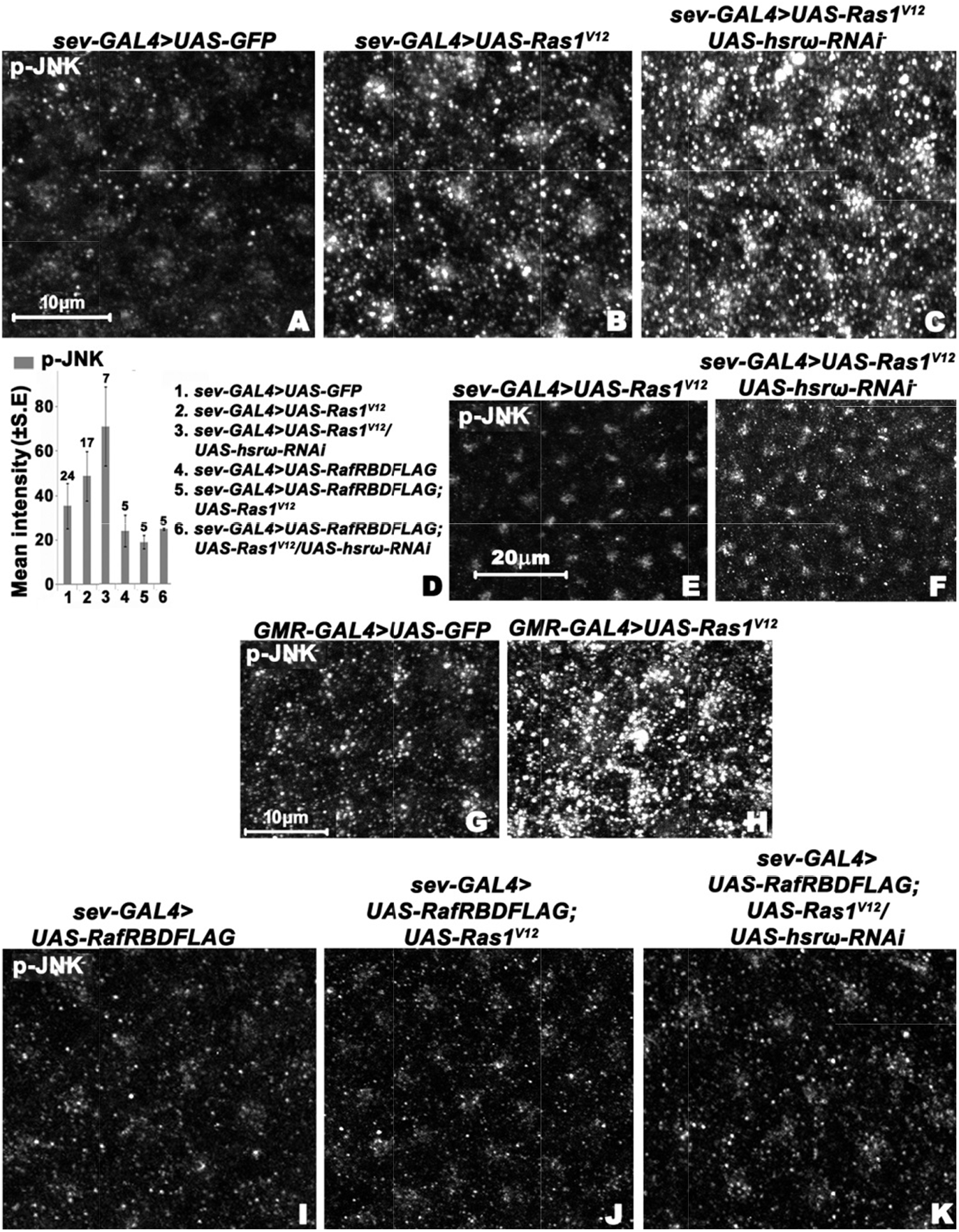
Expression of phosphorylated JNK (pJNK) is enhanced in larval and pupal eye discs of early dying pupae. **A-C** and **E**-K Confocal projection images of third instar larval (**A-C** and **G**-K) and 8-9 Hr pupal (**E**-F) eye discs showing pJNK (white) in different genotypes (noted above the panels). Scale bars in **A**, **G** (10μm) and **E** (20μm) apply to **A-C, G-K** and **E**-F, respectively. **D** Bars showing mean (+ S. E.) fluorescence intensity (Y-axis) of pJNK in the late third instar larval eye discs of different genotypes (1-6, indexed on right side); number on top of each bar indicates the number of eye discs examined.

Similar to *sev-GAL4>UAS-Ras1^V12^ hsrω-RNAi* eye discs, the *GMR-GAL4>UAS-Ras1^V12^* (Fig. 5G) too showed high increase in pJNK levels when compared with *GMR-GAL4>UAS-GFP* control late third instar larval eye discs (Fig. 5H).

It is notable that the pJNK staining increased not only in the *sev-GAL4* expressing cells in ommatidial units of *sev-GAL4>UAS-Ras1^V12^ hsrω-RNAi* eye discs, but also in neighboring non-GFP expressing cells and in the more distant peripodial cells (Fig. 6), which is significant as Dilp8 level too was seen to be increased in these cells in pupal stages (Fig. 4).

**Fig. 6.**
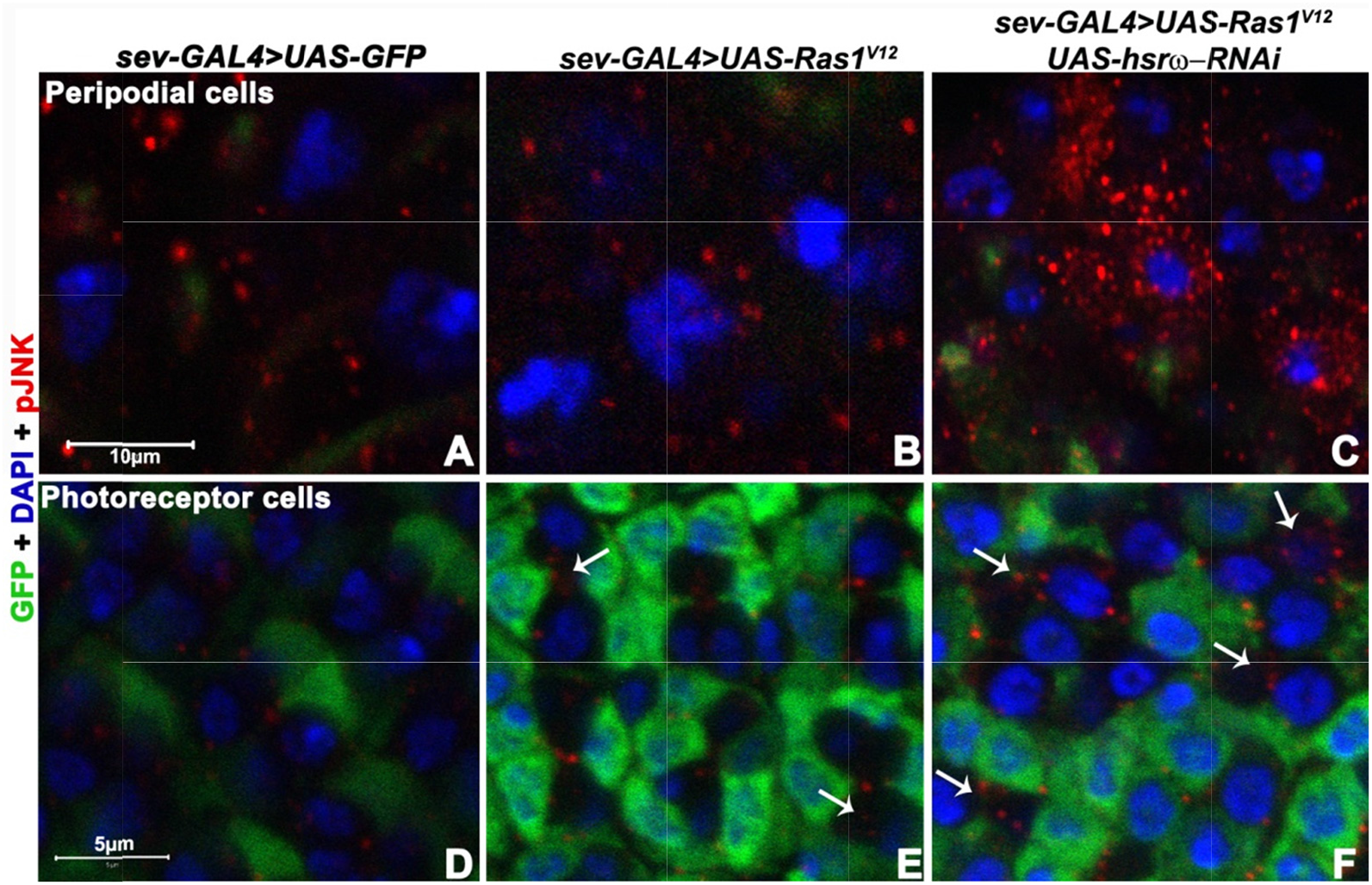
pJNK level is enhanced non-cell-autonomously in eye discs of genotypes that show early pupal death. **A-F** Confocal optical sections of third instar larval eye disc peripodial cells (**A-C**) and of photoreceptor cells (**D-F**) showing pJNK (red) in different genotypes (noted on top of each column), counterstained with DAPI (blue). *sev-GAL4>UAS-GFP* expressing cells are green. White arrows in **E-F** mark the non-GFP expressing cells showing non-cell-autonomous pJNK staining. Scale bars in **A** (10μm) and **E** (5μm) apply to **A-C** and **D-F**, respectively.

### Reduction in Ras signaling in eye discs restored normal pJNK levels in eye discs and rescued the early pupal lethality

We co-expressed *UAS-RafRBDFLAG* in these genotypes to ascertain if the elevated Ras signaling dependent increase in pJNK levels in eye discs co-expressing *sev-GAL4>Ras1^V12^* with *hsrω-RNAi* was indeed responsible for the early pupal death. The RafRBDFLAG acts as a dominant negative suppressor of Ras signaling (Freeman et al., 2010) and thus reduces the damage to eye discs in *sev-GAL4>UAS-Ras1^V12^* and *sev-GAL4>UAS-Ras1^V12^ hsrω-RNAi* larvae (Ray et al., 2019b). Interestingly, there was no pupal lethality when *RafRBDFLAG* was co-expressed with *sev-GAL4>UAS-Ras1^V12^* without or with *hsrω-RNAi* expression (see rows 8-10 in Table 5). Concomitantly, the pJNK levels were also not elevated in *sev-GAL4>UAS-Ras1^V12^* or *sev-GAL4>UAS-Ras1^V12^ hsrω-RNAi* eye discs co-expressing RafRBDFLAG when compared with those in *sev-GAL4>UAS-GFP/UAS-RafRBDFLAG* eye discs (Fig. 5D and Fig. 5I-K).

**Table 5.** High pJNK and Dilp8 in larval and pupal imaginal discs correlate with early pupal lethality

| Genotype |  | pJNK levels in 3rd instar discs (relative to control expressing only *UAS-GFP*) | Dilp8 levels in early pupal discs and brain lobes |
| --- | --- | --- | --- |
| **Eye disc** |  |  |  |
| 1. *GMR-GAL4>UAS-RasI<sup>V12</sup>* | 100% Early pupal death | Very high | High |
| 2. *GMR-GAL4>UAS-rpr* | 100% Early pupal death | Very High | High |
| 3. *GMR-GAL4>UAS-tak1* | 100% Early pupal death | Very high | High |
| 4. *GMR-GAL4>UAS-127Q* | 100% fly emergence | Normal | Normal |
| 5. *sev-GAL4>UAS-GFP* | 100% fly emergence | Normal | Normal |
| 6. *sev-GAL4>UAS-RasI<sup>V12</sup>* | 88% Late pupal death; 12 % fly emergence | Moderately high | Normal |
| 7. *sev-GAL4>UAS-RasI<sup>V12</sup>hsro-RNAi* | 96% Early pupal death; 4% late pupal death (Ray et al., 2019b) | Very high | High |
| 8. *sev-GAL4>RAFRBDFLAG* | 100% fly emergence | Low | Not examined |
| 9. *sev-GAL4>UAS-RasI<sup>V12</sup>RAFRBDFLAG* | 100% fly emergence | Low | Not examined |
| 10. *sev-GAL4>UAS-RasI<sup>V12</sup>RAFRBDFLAG hsro-RNAi* | 100% fly emergence | Low | Not examined |
| **Wing disc** |  |  |  |
| 11. *MS1096-GAL4>UAS-GFP* | 100% fly emergence | Not examined, normal expected | Normal |
| 12. *MS1096-GAL4>UAS-RasI<sup>V12</sup>* | 99% early pupal lethality | Not examined, high expected | High |
| 13. *MS1096-GAL4>UAS-tak1* | 60% early and 25% late pupal lethality | Not examined, high expected | High |
| 14. *MS1096-GAL4>UAS-rpr* | No early pupal lethality but 60% late pupal lethality | Not examined | Slightly elevated |

Shaded cells indicate correlation between early pupal death and very high pJNK and Dilp8 levels in larval and pupal discs, respectively.

In order to further check that the *sev-GAL4* driven elevated Ras expression in eye discs leads to pupal death through elevated JNK signaling and consequent high Dilp8 secretion, we coexpressed *UAS-bsk^DN^*, a dominant negative suppressor of JNK signaling (Martín-Blanco et al 1998 (Martín-Blanco et al., 1998), or *UAS-Dilp8RNAi* (Garelli et al., 2012) in *sev-GAL4>UAS-Ras1^V12^ hsrω-RNAi* eye discs. However, co-expression of neither *UAS-bsk^DN^* nor *UAS-Dilp8RNAi* rescued the early pupal lethality (data not presented). We believe that the ineffectiveness of *UAS-bsk^DN^* or *UAS-Dilp8RNAi* co-expression in suppressing the early pupal lethality associated with *sev-GAL4>UAS-Ras1^V12^ hsrω-RNAi* is related to the earlier noted non-cell-autonomous action of activated Ras in *sev-GAL4* driven *UAS-Ras1^V12^ hsrω-RNAi* eye discs (Ray et al., 2019b) so that pJNK (also see Fig. 6) and Dilp8 (Fig. 4) levels are also enhanced in peripodial and other photoreceptor cells that do not express the *sev-GAL4* driver. On the other hand, action of *sev-GAL4* driven *UAS-bsk^DN^* or *UAS-Dilp8RNAi* remains confined to the *sev-GAL4* expressing cells only. Thus, the *sev-GAL4>UAS-bsk^DN^* or *sev-GAL4>UAS-Dilp8RNAi* expression fails to suppress the high JNK signaling emanating from the other cells in *UAS-Ras1^V12^ hsrω-RNAi* eye discs.

### High JNK and *dilp8* levels in eye or wing discs induced by expression of certain other transgenes also causes early pupal death

We examined if other cases of early pupal lethality following expression of certain transgenes in imaginal discs are also associated with elevated JNK and Dilp8 signaling. We compared pupal lethality, and *dilp8* and/or pJNK levels in *GMR-GAL4>UAS-GFP* or *UAS-Ras1^V12^* with those in *GMR-GAL4> UAS-127Q, GMR-GAL4>UAS-rpr* or *GMR-GAL4> UAS-tak1* eye discs. *GMR-GAL4* driven expression of *UAS-GFP* or *UAS-127Q* does not cause any pupal lethality (Table 5 and also see (Mallik and Lakhotia, 2009b)) while that of *UAS–rpr* (Table 5, also see (Morris et al., 2006)) or *UAS-tak1* (Table 5) caused early pupal death similar to that in *GMR-GAL4>UAS-Ras1^V12^* or *sev-GAL4>UAS-Ras1^V12^ hsrω-RNAi*. Immunostaining for pJNK in late larval eye discs of these different genotypes revealed that *GMR-GAL4>UAS-127Q* (Fig. 7A) showed normal pJNK while those expressing *GMR-GAL4>UAS-rpr* or *UAS-tak1* displayed high pJNK staining (Fig 7B, C). Relative levels of pJNK in different genotypes, determined though quantification of pJNK immunostaining in different genotypes, are shown in Fig. 7D.

**Fig. 7.**
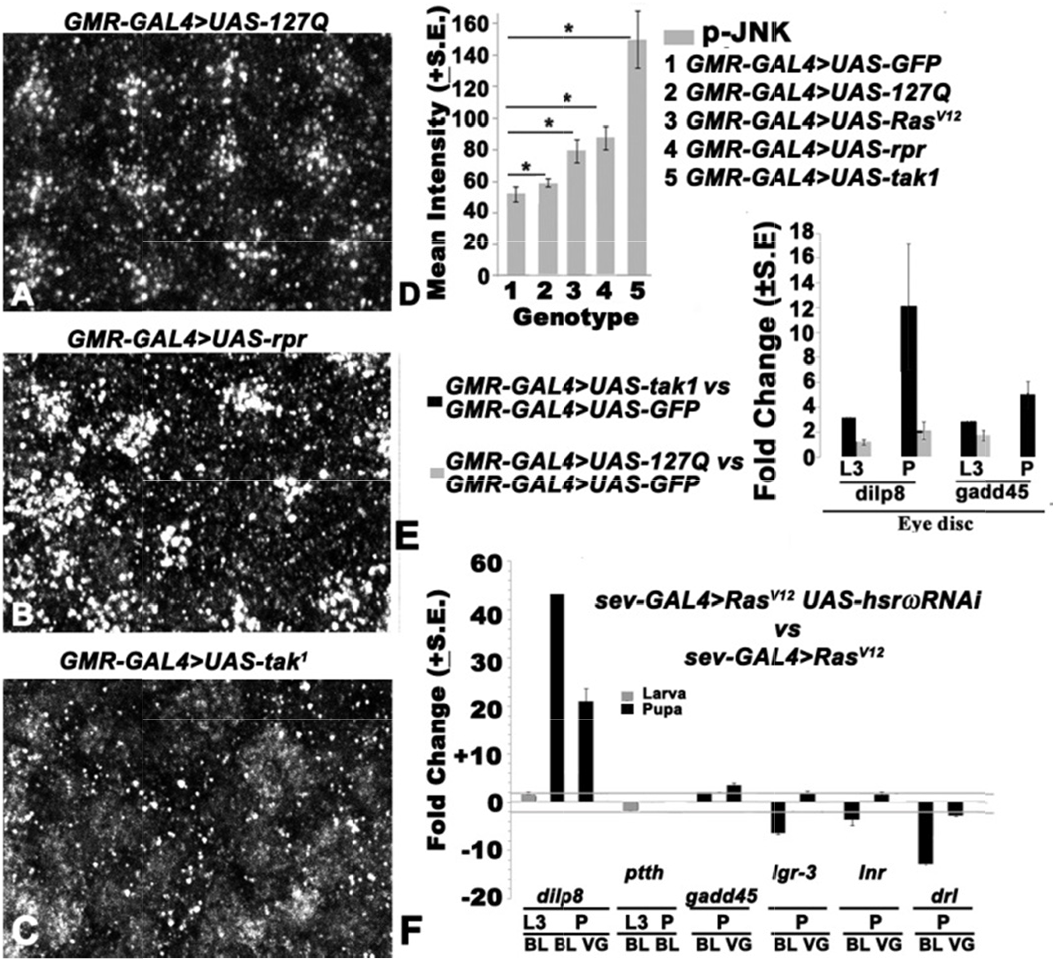
Elevated pJNK levels in eye discs of genotypes showing early pupal death is associated with increased Dilp8 but greatly suppressed *ptth* transcripts in brain ganglia of 8-9 Hr old pupae. **A-C** Confocal projection images of parts of third instar larval eye discs in genotypes mentioned on top of each panel showing distribution of pJNK. Scale bar in **A** denotes 10μm and applies to **A-C**. **D** Bars showing mean (+ S. E., N = 10 discs in each case) fluorescence intensity (Y-axis) of pJNK in the eye discs of different genotypes (1-5, indexed on right); asterisks marks indicate significant (P<0.05) difference in mean values between the pairs of bars connected by horizontal line. **E-F** Bars showing qRT-PCR based mean (+S.E. N = 2 biological replicates) fold changes (Y-axis) in levels of different transcripts (X-axis) in eye discs of *GMR-GAL4>UAS-tak1* (dark gray bars) and *GMR-GAL4>UAS-127Q* (light gray bars) late 3rd instar larvae (L3) or 8-9 Hr old pupae (P) when compared to control *GMR-GAL4> UAS-GFP* (**E**) and in brain lobes (BL) or ventral ganglia (VG) of *sev-GAL4>UAS-Ras1^V12^ hsrω-RNAi* late 3rd instar larvae (L3) or 8-9 Hr old pupae (P) when compared with corresponding *sev-GAL4>UAS-Ras1^V12^* samples. Horizontal grey lines parallel to the X-axis in **F** mark the plus and minus two-fold changes, respectively.

Tak1, a JNKKK, is a known up-regulator of JNK signaling (Weston and Davis, 2002; Silverman et al., 2003; Stronach, 2005; Geuking et al., 2009). When *UAS-tak1* was expressed using a single copy of *sev-GAL4* driver, all progenies (N = 500) eclosed with a bar-like eye. However, driving *UAS-tak1* with two copies of *sev-GAL4*, which would elevate JNK activity much more, caused death of all progeny (N=500) at early pupal stage similar to that after *GMR-GAL4*>*UAS-tak1*, confirming importance of JNK levels in driving the downstream events.

To further confirm enhanced JNK signaling and consequent Dilp8 activity, we measured levels of transcripts of the JNK pathway gene *gadd45*, and *dilp8* in late larval and early pupal *GMR-GAL4>UAS-tak1* and *GMR-GAL4>UAS-127Q* eye discs. In agreement with the absence of pupal death in *GMR-GAL4>UAS-127Q*, neither *dilp8* nor *gadd45* transcripts showed any appreciable change when compared to *GMR-GAL4>UAS-GFP* control. On the other hand, the early dying *GMR-GAL4>UAS-tak1* individuals showed enhanced *dilp8* transcripts at pupal than at larval stages (Fig. 7E). The *gadd45* transcripts also showed significant increase in *GMR-GAL4>UAS-tak1* early pupal eye discs (Fig. 7E).

We examined if increased Ras and JNK activity in wing discs can also cause increased Dilp8 and thus early pupal lethality. We used *MS1096-GAL4* to drive *UAS-Ras1^V12^, UAS-tak1*, or *UAS-rpr* expression in the wing pouch region (Capdevila and Guerrero, 1994). *MS1096-GAL4* driven expression of activated Ras caused near complete (~99%, N=500) early pupal lethality, while that of *UAS-tak1* led to about 60% (N=500) death at early pupal stage; some died as late pupa while only ~15% eclosed with thoracic defects and nearly-absent wings (Table 5). Unlike *GMR-GAL4>UAS-rpr, MS1096-GAL4>UAS-rpr* expression did not cause early pupal death. However, about 60% (N=200) of them died as late pupae while 40% eclosed with vestigial wings. Wing discs of 8-9 Hr old *MS1096-GAL4>UAS-Ras1^V12^* and *MS1096-GAL4 UAS-tak^1^* pupae showed high *dilp8* (Table 5). The *MS1096-GAL4>UAS-rpr* wing discs showed much less increase in *dilp8* transcripts than in *MS1096-GAL4>UAS-Ras1^V12^* or *MS1096-GAL4 UAS-tak^1^* (Table 5), which correlates with survival of many of them through the early pupal period.

A summary of pupal survival, pJNK and Dilp8 levels in the different genotypes examined in our study is presented in Table 5, which shows that both the JNK and Dilp8 signaling cascades were up-regulated in larval and early pupal eye discs, respectively, in all the genotypes that displayed high death at about 23-25 Hr APF. The increased Ras signaling in the eye disc seem to elevate JNK which induces synthesis of Dilp8 in eye or wing discs.

### Elevated Ras signaling in eye discs leading to lowered Ptth level in brain seems to be responsible for lowered ecdysone and subsequent early pupal death

Since our earlier report (Ray and Lakhotia, 2015) showed that both the *sev-GAL4* and *GMR-GAL4* drivers also express in the seven segmentally arranged paired dorso-medial neurons in larval and pupal ventral ganglia (VG) and certain other neurons in brain, we checked if the early pupal death was related to ectopic expression of activated Ras in these VG neurons.

We compared levels of *dilp8, ptth, gadd45, lgr3, inr* and *drl* transcripts in brain lobes and ventral ganglia of 8-9 Hr old pupae *sev-GAL4>UAS-Ras1^V12^hsrω-RNAi* and *sev-GAL4>UAS-Ras1^V12^* using qRT-PCR (Fig. 7F). Although *dilp8* transcripts showed elevation in the *sev-GAL4>UAS-Ras1^V12^ hsrω-RNAi* in pupal, but not larval, brain, the JNK pathway gene *gadd45* did not show any significant change in pupal brain (Fig 7F). The *ptth* transcript levels were slightly reduced in larval brain, but were nearly absent in pupal brain of *sev-GAL4>UAS-Ras1^V12^ hsrω-RNAi* individuals (Fig 7F). This is significant since Ptth hormone from brain triggers the prothoracic gland to produce ecdysone (Rewitz et al., 2009; Niwa and Niwa, 2016) and thus such low level of *ptth* transcripts (Fig 7F) indicates absence of this triggering mechanism in individuals with high Ras activity in the eye discs.

The *lgr3, inr* and *drl* transcripts showed somewhat reduced levels in *sev-GAL4>UAS-Ras1^V12^ hsrω-RNAi* brain lobes but little change in ventral ganglia of 8-9 Hr *sev-GAL4>UAS-Ras1^V12^ hsrω-RNAi* pupae (Fig. 7F).

Immunostaining of 8-9 Hr old pupal brain ganglia for pJNK revealed that its expression was not affected by *sevGAL4>UAS-Ras^V12^* expression, without or with concurrent down-regulated *hsrω* levels. Brain lobes showed comparable JNK staining, with a stronger expression in the Mushroom body, in all the genotypes (Fig. 8A-C).

**Fig. 8.**
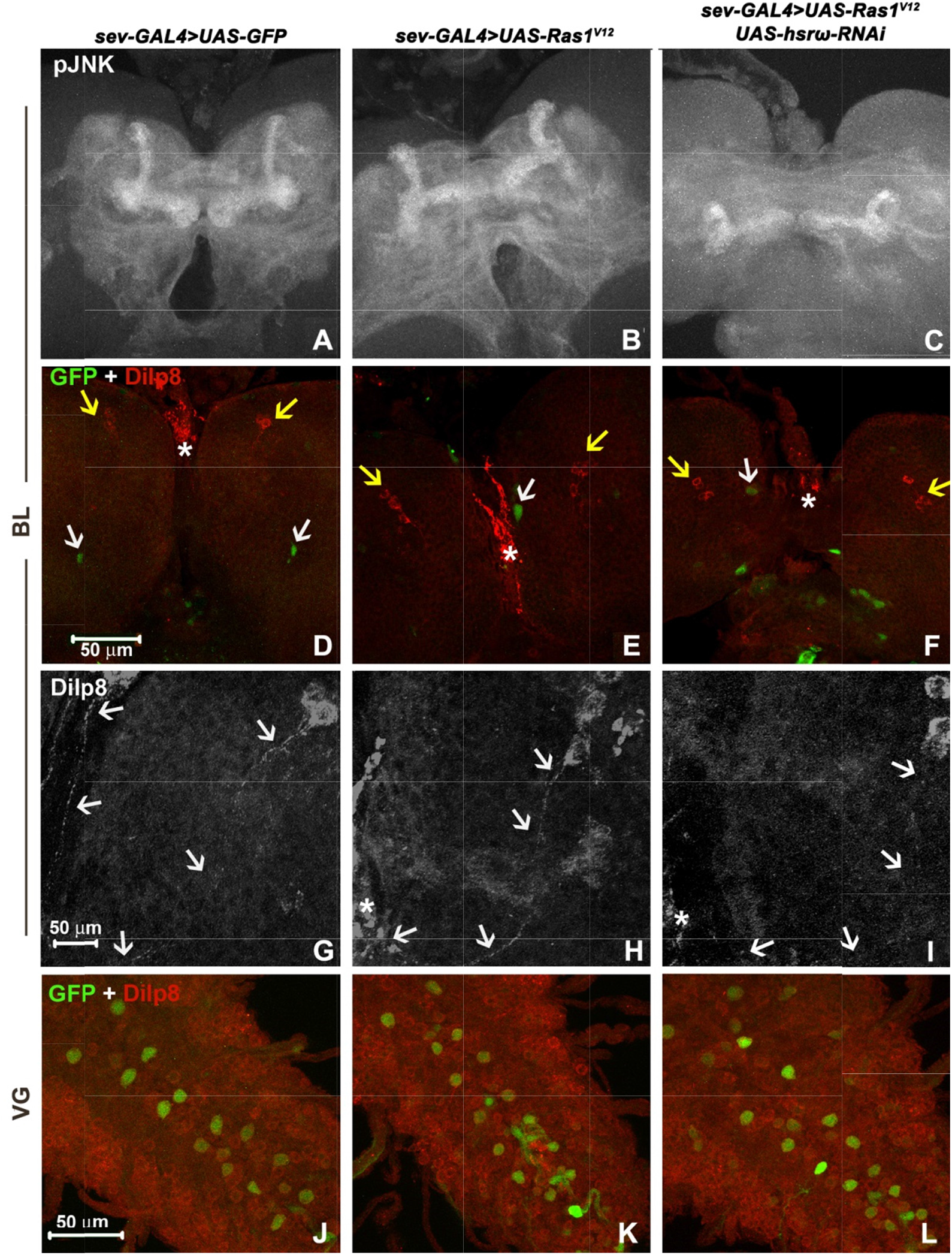
Expression of *sev-GAL4>UAS-Ras1^V12^*, without or with co-expression of *hsrω-RNAi*, in eye discs and specific neurons in the central nervous system (CNS) does not increase pJNK and Dilp8 levels in the CNS. **A-L** Confocal projection images of parts of brain lobes (**A-I**) and ventral ganglia (**J-L**) of 8-9 Hr old pupae of different genotypes (noted on top of each column) showing pJNK (white in **A-C**) and Dilp8 (red in **D-F** and **J-L**, white in **G-I**) and *sev-GAL4>GFP* (green in **D-F** and **J-L**). White arrows in D-F indicate the GFP-positive neurons in antero-medial part while the yellow arrows indicate the mid-lateral groups of four Dilp8 expressing neurons in each brain lobe; the basal region of prothoracic gland is marked with asterisk. **G-I** show higher magnification images of Dilp8 staining in one of the brain lobes with the white arrows indicating the Dilp8 positive axonal projections from the group of Dilp8 positive neurons to the basal region of prothoracic gland (marked by asterisk). The scale bars in **D** (50 μm), **G** (20 μm) and **J** (50 μm) apply to **A-F, G-I** and **J-L**, respectively.

A low abundance of Dilp8 was seen all through the brain and ventral ganglia of *sevGAL4>UAS-GFP* (Fig. 8D, G, J). Interestingly, none of the *sevGAL4>UAS-GFP* and *Ras1^V12^* transgene expressing neurons in brain and ventral ganglia (Ray and Lakhotia, 2015) of *sev-GAL4>UAS-Ras1^V12^*, showed elevation in Dilp8 expression beyond the low expression seen in adjoining cells (Fig. 8E, H, K). Likewise, the Dilp8 expression in the 8-9 Hr old pupal *sev-GAL4>UAS-Ras1^V12^ hsrω-RNAi* brain lobes and ventral ganglia remained similar to that in *sevGAL4>UAS-GFP* and *sev-GAL4>UAS-Ras1^V12^* (Fig. 8F, I, L). Intriguingly, a group of 4 neurons in mid-lateral part in each of the brain-hemispheres *in all the four genotypes* showed strong Dilp8 presence in cytoplasm and their axonal projections, which followed a semi-circular path to reach posterior part of the prothoracic gland (Fig. 8D-I). The posterior part of prothoracic gland also showed strong Dilp8 staining. These might be responsible transmitting Dilp8 mediating Ptth signaling to prothoracic glands.

However, contrary the above noted increase in *dilp8* transcripts in early pupal CNS of *sev-GAL4>UAS-Ras1^V12^ hsrω-RNAi*, Dilp8 protein staining in CNS, including in the specific 4 pairs of neurons and the basal part of the prothoracic gland, remained comparable in all the genotypes. Thus, despite the increase in *dilp8* transcripts, the *sev-GAL4* driven expression of activated Ras without or with concurrent altered *hsrω* expression did not detectably affect Dilp8 protein levels in the CNS.

## Discussion

A major objective of the present study was to understand reasons for early pupal death following expression of certain transgenes driven by the predominantly eye-specific *sev-GAL4* or *GMR-GAL4* drivers (Morris et al., 2006; Mallik and Lakhotia, 2009b; Ray et al., 2019b). It is known that besides the eye disc, the *sev-GAL4* as well as *GMR-GAL4* drivers also express in specific but common sets of neurons in the central nervous system (Ray and Lakhotia, 2015). While the two drivers express nearly equally in the BG and VG, the *GMR-GAL4* is expressed much more extensively in eye discs, covering nearly all cells posterior to the morphogenetic furrow, the *sev-GAL4* expression is limited to the R7 lineage of cells (Ray and Lakhotia, 2015). As expected the ectopic Ras activity was much higher in *GMR-GAL4>UAS-Ras1^V12^* eye discs than in *sev-GAL4>UAS-Ras1^V12^* discs and this was paralleled by early pupal death when *Ras1^V12^* was expression was driven by the *GMR-GAL4* (Ray et al., 2019b). The early pupal death of *sev-GAL4>UAS-Ras1^V12^ UAS-hsrω-RNAi* individuals was also associated with much higher levels of activated Ras (Ray et al., 2019b). Our finding that expression of *Ras1^V12^* or certain other transgenes in wing imaginal discs under the *MS1096-GAL4* driver, which is not known to be expressed in central nervous system, too was associated with early pupal death, provides further support for the view that the initial signal that triggers early pupal death emanates from imaginal discs rather than from the BG or VG. This view is further supported by the absence of any difference in pJNK or Dilp8 activity in BG or VG of early dying and normally surviving early pupae.

As reported earlier (Colombani et al., 2012; Garelli et al., 2012; Okamoto and Yamanaka, 2015), our results also suggest Dilp8 to be the messenger between eye discs and prothoracic gland via brain to affect ecdysone biosynthesis. Previous studies established a role of Dilp8 in regulating larval development by delaying ecdysone synthesis when imaginal disc development is adversely affected either because of tumor formation or other kind of damage, e.g., expression of apoptotic genes, or inadequate growth (Colombani et al., 2012; Garelli et al., 2012; Colombani et al., 2015; Garelli et al., 2015; Okamoto and Yamanaka, 2015). Present results show that highly elevated ectopic Ras signaling in eye discs also induces Dilp8 expression. However, rather than affecting the late larval ecdysone release, in this case the post-pupation ecdysone surge is inhibited. Our results show that a very high increase in Ras signaling in larval eye or wing discs (*GMR-GAL4>UAS-Ras1^V12^, sev-GAL4>UAS-Ras1^V12^ hsrω-RNAi, MS1096-GAL4>UAS-Ras1^V12^*) caused significant elevation of Dilp8 during early pupal life (>8 Hr APF) which was associated with great reduction in *ptth* transcripts in the brain and reduced activity of ecdysone biosynthesis genes in prothoracic gland. Consequently, the ecdysone surge that normally occurs after 8 Hr pupa formation (Handler, 1982) could not occur. Our study thus reveals that the ecdysone release after 8 Hr pupa formation is another check point where the organism can halt further development when some organ is not normally functioning or developing. Our finding that the early pupal death following ectopic expression of several other transgenes like *UAS-tak1, UAS-rpr* etc under eye- or wing-disc specific GAL4 drivers too is associated with elevated Dilp8 levels suggests Dilp8 to be a common and essential inter-organ messenger between epithelia and the brain during larval as well as pupal metamorphosis.

It is believed that damaged or relatively poorly growing larval imaginal discs show high JNK signaling which leads to secretion of Dilp8 to delay pupation (Andersen et al., 2015; Katsuyama et al., 2015). Our results also showed elevated transcripts of several JNK pathway members and pJNK activity in *GMR-GAL4>UAS-Ras1^V12^* and *sev-GAL4>UAS-Ras1^V12^ hsrω-RNAi* eye discs. Likewise, we found elevated JNK and Dilp8 signaling following expression of other transgenes in eye or wing discs that also cause early pupal lethality. Further support for the pJNK’s role in elevation of Dilp8 levels derives from prevention of high JNK and Dilp8 levels, and early pupal lethality following co-expression of RafRBDFLAG, which acts as a dominant negative suppressor of Ras signaling (Freeman et al., 2010). The observed lack of any rescue in early pupal lethality following co-expression of *UAS-bsk^DN^* or *UAS-Dilp8RNAi* with *sev-GAL4>UAS-Ras1^V12^ hsrω-RNAi* does not contradict the role of JNK and Dilp8 in the observed early pupal lethality since unlike the non-cell-autonomous action of activated Ras in eye discs (Ray et al., 2019b), action of the *UAS-bsk^DN^* or the *UAS-Dilp8RNAi* remains limited to the cells in which the *sev-GAL4* driver is expressed. Consequently, the down-regulation of JNK or Dilp8 expression in only the *sev-GAL4* expressing cells does not suppress the spread of active Ras to neighbouring cells where it triggers elevation in JNK and Dilp8 signaling. On the other hand, expression of RafRBDFLAG not only inhibits Ras activity in the *sev-GAL4* expressing source cells but also in the neighbouring cells since the active Ras moves non-autonomously together with the RafRBDFLAG (Ray et al., 2019b) so that the wider Ras signaling and consequent pupal death are completely suppressed.

It is interesting that although very high Ras signaling was associated with increased pJNK and Dilp8 levels in third instar larval eye discs of early dying pupae, ecdysone biosynthesis pathway genes in prothoracic gland were not affected and the third instar larvae pupated normally. This situation is different from the earlier reported cases of delayed pupation in larvae with damaged/abnormally developing imaginal discs (Colombani et al., 2012; Garelli et al., 2012; Andersen et al., 2015; Colombani et al., 2015; Garelli et al., 2015; Okamoto and Yamanaka, 2015). The normal pupation, despite high JNK levels in the larval imaginal discs in *GMR-GAL4>UAS-Ras1^V12^* or *sev-GAL4>UAS-Ras1^V12^ hsrω-RNAi* larvae is possibly due to a sub-threshold elevation in Dilp8 levels in late larval stages. Apparently, the small increase in Dilp8 signaling failed to significantly down-regulate Ptth activity so that ecdysone biosynthesis continued in larval prothoracic glands, and, therefore, the pupation was neither prevented nor significantly delayed. However, as the very high Ras and JNK signaling continued in early pupal imaginal discs, the Dilp8 levels increased steeply beyond the threshold levels and substantially inhibited Ptth synthesis in brain and thus reduced ecdysone biosynthesis in prothoracic glands in >8 Hr old pupae. In absence of the ecdysone surge in >8 Hr old pupae (Handler, 1982), the next set of metamorphic changes failed to occur. Thus, our study besides confirming the inter-tissue signaling role of Dilp8 in maintaining developmental homeostasis, reveals existence of a novel Dilp8-mediated checkpoint. We show that onset of early pupal metamorphosis is also stage at which Dilp8 can act. Apparently, while the larval duration can be extended by delaying pupation till the organism attains homeostasis between different organs, the pupal development does not have such a cushion so that in the absence of the required ecdysone trigger, the pupae die without completing their initial metamorphic changes. It would be interesting to examine if the late pupal/pharate stage lethality observed in many genotypes is also related to Dilp8 activity.

Our results provide further evidence for two interesting phenomena related to developmental dynamics. First, changes in local tissue environment cause systematic changes with consequences at organism level, and second, certain developmental stages are more susceptible to such effects. Present findings showing ectopic high Ras signaling as a trigger pathway for Dilp8 synthesis in epithelial cells raise the possibility that ectopic expression of activated Ras in certain diseases/tumors may also have far reaching organism level consequences through down-stream signaling cascades.

## Acknowledgements

We acknowledge the Bloomington *Drosophila* Stock Center (USA) for fly stocks, and Dr. P. Leopold (France) for the anti-Dilp8. We thank the DBT-BHU Interdisciplinary School of Life Sciences for the Microarray and real time PCR facilities. We also thank the Centre of Advanced Studies in Department of Zoology, BHU for various facilities.

## Author contributions

MR and SCL planned experiments, analyzed results and wrote the manuscript. MR carried out the experimental work and collected data.

## Data availability

The microarray data have been deposited at GEO repository (http://www.ncbi.nlm.nih.gov/geo/query/acc.cgi?acc=GSE80703).

